# Reconciling intrinsic and ecosystem water-use efficiency reveals physiological mechanisms underlying carbon–water coupling across biological scales

**DOI:** 10.64898/2026.10.07.757244

**Authors:** Paulina F. Puchi, Daniela Dalmonech, Daniele Castagneri, Elia Vangi, Marco M. Lehmann, Matthias Saurer, Stefan Klesse, Lukas Hörtnagl, Alessio Collalti

## Abstract

Understanding how carbon–water coupling scales from leaves to ecosystems remains a major challenge because water-use efficiency (WUE) metrics integrate physiological, hydrological, and ecological processes. We integrated tree-ring width, triple stable isotopes (δ¹³C, δ^18^O and δ²H), eddy covariance (EC) observations, and a process-based model to investigate the drivers of intrinsic (_i_WUE) and ecosystem water-use efficiency (WUE_eco_) across biological scales in a mature *Fagus sylvatica* forest between 2004 and 2023. Modelled and isotope-derived _i_WUE were strongly correlated (r = 0.61, *p* < 0.05) and both indicated an approximately 20% long-term increase, consistent with increasing gross primary productivity. In contrast, WUE_eco_ derived from EC was lower than model estimates and exhibited a positive trend not reproduced by the model. This divergence primarily reflected differences in biological representation and hydrological integration, as the single-species model underestimated ecosystem evapotranspiration by excluding co-occurring deciduous and evergreen species together with understory vegetation. Triple-isotope analyses showed that the long-term increase in _i_WUE was driven by enhanced photosynthetic assimilation rather than sustained stomatal closure, whereas δ^18^O and δ²H variability was mainly controlled by source-water dynamics. These findings provide a mechanistic framework for reconciling leaf- and ecosystem-scale WUE and improving predictions of forest carbon–water coupling under climate change.

## Introduction

Understanding how forests coordinate carbon uptake and water loss is central to predicting terrestrial ecosystem responses to climate change. Carbon assimilation through photosynthesis and water loss through transpiration are tightly coupled via stomatal regulation, making water-use efficiency (WUE) a key integrative metric of plant physiological functioning. WUE describes the balance between carbon gain and water loss and therefore reflects how vegetation optimizes resource use under changing environmental conditions (Farquhar et al., 1982; Medlyn et al., 2017). At regional and global scales, shifts in WUE can strongly influence carbon sequestration, evapotranspiration (ET), and land–atmosphere feedbacks, making this parameter critical for constraining Earth System Models and forecasting ecosystem responses to rising atmospheric CO_2_ and intensifying hydroclimatic stress (Keenan et al., 2013; Peters et al., 2018).

However, WUE is not a scale-invariant property, and its interpretation depends strongly on the spatial and temporal scale at which it is measured (Beer et al., 2009). Indeed, although WUE is widely adopted within the scientific community as a key indicator of plant performance under water-limited conditions, its meaning has progressively broadened over time. As a result, WUE is now characterized by multiple, coexisting definitions, each tailored to specific methodological approaches or research contexts. While this conceptual flexibility has facilitated its application across diverse disciplines, it has also led to a certain degree of ambiguity, ultimately limiting the comparability of results across studies and contexts (Puchi et al., 2025). At the leaf level, intrinsic water-use efficiency (_i_WUE) is defined as the ratio between net photosynthetic assimilation (A) and stomatal conductance to water vapor (*g_s_*). Tree-ring δ^13^C provides a retrospective archive of _i_WUE because carbon isotope discrimination during photosynthesis reflects the ratio of intercellular to atmospheric CO_2_ concentrations (cᵢ/cₐ), which is controlled by the balance between assimilation and stomatal conductance (Farquhar et al., 1982). Consequently, isotope-derived _i_WUE integrates physiological regulation over the period of wood formation and provides a powerful tool to reconstruct long-term plant responses to environmental variability and atmospheric CO_2_ increase (Saurer & Voelker, 2022).

In contrast, ecosystem-scale WUE (WUE_eco_) is typically quantified using eddy covariance measurements as the ratio between gross primary productivity (GPP) and evapotranspiration. This flux-based metric integrates canopy-level carbon uptake and total ecosystem water losses, including transpiration, soil evaporation, and canopy interception (Knauer et al., 2018). As such, WUE_eco_ represents an emergent property of vegetation structure, atmospheric demand, soil hydrology, and land–atmosphere coupling. Critically, the inclusion of non-transpirational water fluxes in ET means that WUE_eco_ does not only reflect stomatal-level physiological regulation, potentially leading to systematic biases when comparing with leaf-level metrics (Dekker et al., 2016; Wang et al., 2020; see Puchi et al. (2025) for a broad overview of the many definitions and equations used).

Bridging leaf-level and ecosystem-level representations of WUE, therefore, remains a major challenge in ecohydrology and ecosystem modeling. Several fundamental issues complicate cross-scale comparisons: (1) differences in temporal integration of tree rings record seasonal to annual signals, while eddy covariance provides sub-daily to daily measurements; (2) spatial representativeness individual trees vs. ecosystem-level fluxes; and (3) hydrological partitioning stomatal transpiration vs. total evapotranspiration (Medlyn et al., 2017). Several studies have reported weak or inconsistent relationships between long-term trends in _i_WUE inferred from tree rings and WUE_eco_ derived from eddy covariance measurements (Seibt et al., 2008; Gea-Izquierdo et al., 2015; Lavergne et al., 2019; Li et al., 2022). These discrepancies highlight that processes measured at the stomatal scale are not directly comparable to those emerging at the ecosystem scale, where non-stomatal water fluxes and canopy structural effects become increasingly important.

Therefore, while partitioning ET into transpiration and evaporation components could theoretically improve cross-scale comparisons, such partitioning remains challenging and uncertain, particularly in complex forest ecosystems (Nelson et al., 2018). Therefore, understanding the relationship between leaf-level _i_WUE and ecosystem-level WUE calculated from total ET remains a fundamental challenge for interpreting tree-ring isotope records in the context of ecosystem functioning.

Additionally, increases in _i_WUE do not always correspond to higher forest growth or productivity, because the relationship between WUE and carbon allocation depends strongly on hydroclimatic constraints such as vapor pressure deficit (VPD), soil moisture availability, and plant hydraulic regulation (Frank et al., 2015; Zhang et al., 2025). Among these factors, VPD plays a central role in linking leaf- and ecosystem-scale WUE because it simultaneously influences stomatal conductance and atmospheric evaporative demand. As VPD increases, transpiration may continue to rise despite partial stomatal closure owing to the steeper leaf-to-air vapor pressure gradient, potentially decoupling leaf-level physiological responses from ecosystem-scale carbon–water exchange (Grossiord et al., 2020; Diao et al., 2024; Medlyn et al., 2017). Consequently, understanding whether long-term changes in _i_WUE are driven primarily by changes in g_s_ or A is critical for interpreting isotope records in the context of ecosystem functioning (Guerrieri et al., 2019). Because δ^13^C reflects changes in cᵢ/cₐ rather than A or g_s_ independently, additional constraints are often required to distinguish whether observed _i_WUE trends arise from shifts in photosynthetic capacity or stomatal regulation (Scheidegger et al., 2000). In particular, oxygen isotopes (δ^18^O), which are primarily sensitive to transpiration and evaporative enrichment of leaf water (Roden et al., 2000; Barbour, 2007), have been combined with δ^13^C records within the conceptual dual-isotope framework proposed by Scheidegger et al. (2000) to attribute _i_WUE changes to either stomatal or biochemical drivers. More recently, hydrogen isotopes (δ²H) have also been explored as complementary tracers of leaf water enrichment and source-water signals, providing additional constraints on plant water relations and ecohydrological processes that can further refine interpretations of stomatal regulation and transpiration dynamics (Lehmann et al., 2021; Saurer et al., 2025). This approach has since been applied in dendroecological studies to interpret long-term isotope trends under changing atmospheric and climatic conditions (e.g., Saurer et al., 2004; Lavergne et al., 2020).

Process-based ecosystem models, such as 3D-CMCC-FEM, provide a promising framework for bridging leaf-level and ecosystem-level WUE estimates because they explicitly simulate canopy photosynthesis, stomatal conductance, transpiration, carbon allocation, respiration, and water balance using mechanistic eco-physiological principles (Fatichi et al., 2016; Keenan et al., 2013; Collalti et al., 2016; 2024; Saponaro et al., 2026). By systematically comparing model simulations with both tree-ring isotope records and eddy covariance observations, we can assess the extent to which models realistically represent the physiological processes driving WUE variability across spatial and temporal scales, while also using models to bridge observational gaps by inferring processes that are difficult or impossible to measure directly. This integrative approach also enables the identification of key mechanisms underlying discrepancies between leaf-level and ecosystem-level estimates, thereby helping to clarify scale-dependent dynamics and improve model performance.

Recent studies have increasingly adopted multi-scale approaches to investigate the responses of WUE to environmental stress. For example, Rambal et al. (2025) combined stable isotopes, gas exchange, and eddy covariance measurements to assess drought effects on intrinsic and ecosystem water-use efficiency in a Mediterranean evergreen oak forest, highlighting the importance of considering biological scale when interpreting carbon–water relationships. However, despite these advances, studies integrating physiological proxies, ecosystem observations, and mechanistic modelling within a unified framework remain lacking.

In this study, we integrate eddy covariance observations, tree-ring stable isotopes (δ^13^C, δ^18^O, δ²H), dendrochronological measurements, and simulations from a process-based model to investigate carbon–water interactions in a temperate mixed forest dominated by *Fagus sylvatica* L. at the Lägeren site in Switzerland over a period of two decades (2004-2023). Specifically, this study addresses three main objectives:

1. *Temporal scaling and integration*: To evaluate whether the agreement between isotope-derived intrinsic _i_WUE (i.e. A/g_s_) and ecosystem WUE_eco_ (i.e. GPP/ET) improves with temporal aggregation (daily → monthly → annual), and to assess whether restricting analyses to the growing season enhances cross-scale comparability (see Box 1);

**Box 1.**
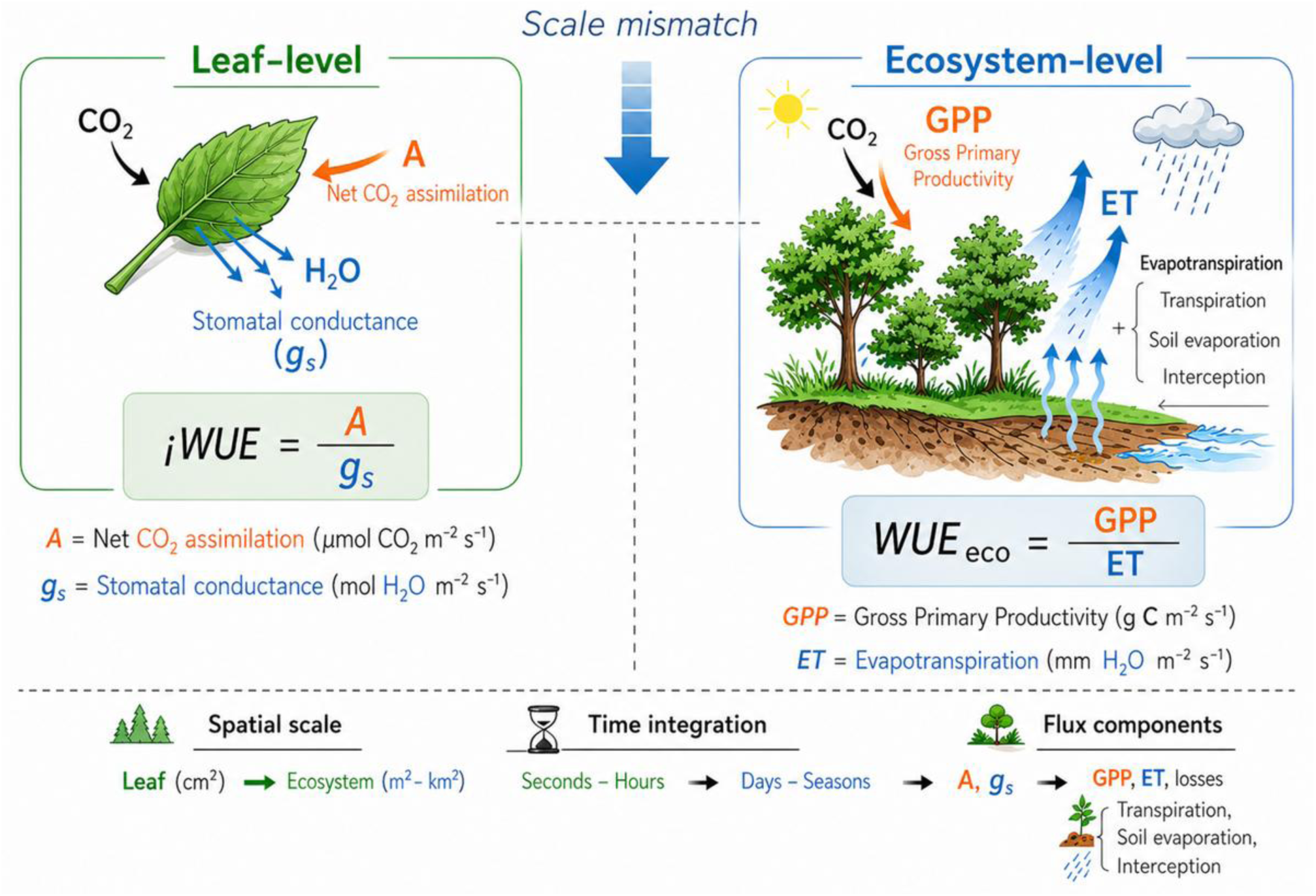
Conceptual framework illustrating the scale mismatch between intrinsic water-use efficiency (ᵢWUE) at the leaf level and ecosystem water-use efficiency (WUE_eco_). At the leaf scale, ᵢWUE is defined as the ratio between net CO_2_ assimilation (*A*) and stomatal conductance to water vapor (*g_s_*), reflecting the physiological trade-off between carbon gain and water loss regulated by stomata. At the ecosystem scale, WUE_eco_ is defined as the ratio of gross primary productivity (GPP) to evapotranspiration (ET), integrating canopy-level carbon uptake and total water fluxes, including transpiration, soil evaporation, and interception losses. The schematic highlights the conceptual and process-based differences between scales, emphasizing that ᵢWUE and WUE_eco_ are governed by distinct flux components, integration times, and spatial domains.
2. *Model-data integration for bridging scales*: To use a process-based model as a mechanistic bridge between leaf-level and ecosystem-level WUE by: (i) validating model-simulated GPP and ET against eddy covariance observations, (ii) comparing model-derived _i_WUE^mod^ with tree-ring isotope-derived _i_WUE^obs^, and (iii) evaluating whether the model can reproduce observed patterns in both leaf-level and ecosystem-level WUE;
3. *Disentangling physiological controls*: To assess the relative contributions of stomatal conductance and photosynthetic assimilation in driving interannual variability in intrinsic _i_WUE, using simulations from the process-based model together with the conceptual dual-isotope framework (Scheidegger et al., 2000). Specifically, the model was used to simulate physiological variables (e.g., stomatal conductance), while the combined interpretation of tree-ring δ^13^C and δ^18^O was used to infer whether changes in _i_WUE were primarily driven by stomatal regulation or by changes in photosynthetic capacity.

By integrating complementary methodologies operating at different levels of ecological organization, from tree-ring stable isotopes and dendrochronology at the leaf level to eddy covariance observations and process-based ecosystem modelling at the ecosystem level, this study presents, to our knowledge, the first mechanistic framework for disentangling WUE across biological scales. Bridging physiological and ecosystem processes may prove essential for advancing our understanding of carbon–water coupling in a changing climate.

## Materials and methods

### Study site and climate

The study was conducted in a managed deciduous mountain forest located on the steep slopes of the Lägeren mountain (northwest of Zurich, Swiss Plateau), at approximately 700 m asl. The forest within the tower footprint is characterized by a mixed-species deciduous stand. The most abundant tree species are the European beech (*Fagus sylvatica* L.), ash (*Fraxinus excelsior* L.), and Norway spruce *(Picea abies* (L.) H.Karst., 1881) (Shekhar et al., 2024). In spring and early summer, bear’s garlic (*Allium ursinum* (L., 1753)) forms a dense understory (Etzold et al., 2011). The site, known as CH-Lae in the global FLUXNET network, has been continuously monitored for carbon, water, and energy fluxes using the eddy covariance technique since April 2004. The tower is located at 47°28′42.0″ N, 8°21′51.8″ E, at an elevation of 689 m a.s.l. (Fig. 1a).

**Fig. 1.**
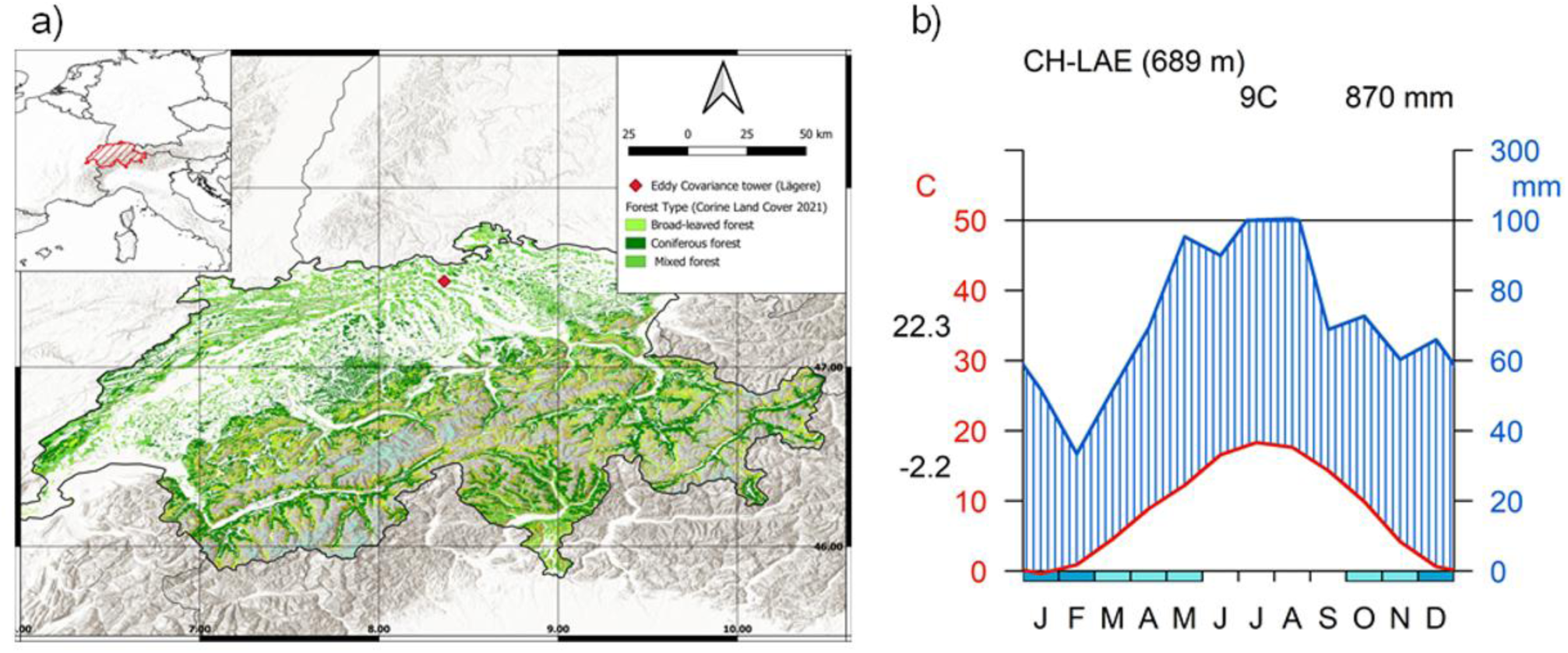
(a) Location of Lägeren CH-Lae site, Canton of Aargau, Switzerland, showing forest cover types from CORINE Land Cover (2021). The red dot indicates the position of the eddy covariance (EC) tower; (b) Walter-Lieth climate diagram for the period 2004-2023, derived from meteorological data recorded at the EC tower. Teal boxes (bottom) are frost indicators: lighter boxes mark months with *probable frost* (mean minimum temperature approaches 0°C); darker boxes mark months with *certain frost* (mean minimum temperature below 0°C). In this diagram, frost risk is concentrated in J, F, M, A, N, D, consistent with the recorded minimum of −2.2°C.

The site is underlain by limestone bedrock, and the main soil types are Rendzic Leptosols (Rendzinas) and Haplic Cambisols, according to the World Reference Base for Soil Resources (IUSS Working Group WRB, 2007). Soils are well-drained, and the litter layer is generally thin, as leaf litter decomposes almost completely within a year due to the relatively mild climatic conditions and high biological activity.

The CH-Lae site is characterized by a temperate humid climate with mild temperatures and abundant precipitation throughout the year. Based on data recorded from 2004 to 2023, the mean annual air temperature is 8.7 °C, and the mean annual precipitation totals approximately 870 mm. Precipitation is relatively evenly distributed across months, with slightly wetter conditions during late spring and summer. The warmest month is July, with a mean temperature of about 22 °C, while the coldest month is January, averaging around –2 °C. Snowfall occurs occasionally in winter but is generally short-lived due to moderate temperatures (Fig. 1b).

### 2.2 Eddy covariance flux measurements

Half-hourly fluxes of energy, water vapour and carbon dioxide (CO_2_) have been continuously measured using the eddy covariance (EC) method since April 2004 (Etzold et al., 2010). The EC system consisted of a three-dimensional sonic anemometer (Gill Solent HS, Gill Instruments Ltd., Lymington, UK) for the measurements of high-frequency (20Hz) wind data, and an open-path (model LI-7500, LI-COR Biosciences, Lincoln, NE, USA; installed between 2004 and 2015) or enclosed-path (LI-7200, from 2016 onwards) infrared gas analyzer (IRGA). The EC instruments were mounted at a height of 47 m above ground, corresponding to approximately 17 m above the mean canopy height. Fluxes were calculated following established community guidelines (Aubinet et al., 2012; Sabbatini et al., 2018). Subsequent data quality control, gap-filling, and flux partitioning into gross primary production (GPP) and ecosystem respiration (R_eco_) followed the standard FLUXNET approach (Pastorello et al., 2020). Detailed site information is available from the Swiss FluxNet network (https://www.swissfluxnet.ethz.ch, site CH-Lae).

Evapotranspiration (ET) was derived from latent heat flux (LE) measurements obtained from the EC system. Half-hourly LE values (W m⁻²) were converted to ET using the latent heat of vaporization of water (λ = 2.45 × 10⁶ J kg⁻¹) according to:

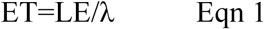

where ET is expressed in kg m⁻² s⁻¹, equivalent to mm s⁻¹. Half-hourly ET estimates were integrated and aggregated to obtain daily ET totals (mm day⁻¹).

The upper platform of the tower is equipped with instruments that record standard meteorological variables, including net radiation (Kipp & Zonen CNR1, Delft, Netherlands), air temperature, and relative humidity (Rotronic MP101A, Bassersdorf, Switzerland). In the vicinity of the flux tower, soil temperature and moisture are monitored at a depth of 5 cm using ECH_2_O EC-20 probes (Decagon Devices, Pullman, WA, USA). Soil heat flux is measured with two HFP01 heat flux plates (Hukseflux, Delft, Netherlands) installed side by side at 3 cm depth. Mean soil heat flux is corrected for heat storage changes in the layer above the plates following the procedure described by Monteith and Unsworth (1990). All measurements are automatically controlled and recorded by a CR10X data logger (Campbell Scientific Inc., Loughborough, UK).

### 2.3 Field sampling and processing dendrochronological data

In July 2024, 15 trees of *Fagus sylvatica* were sampled within the eddy covariance tower footprint, targeting the upslope sector of the stand where beech dominance is highest (approximately 85% of basal area). Cores were extracted at 1.3 m from the ground using a 5 mm increment borer, with two cores extracted per tree. In the tree-ring laboratory of the Federal Swiss Institute for Forest, Snow, and Landscape Research (WSL), wood cores were air-dried and polished with sandpaper of successively increasing grains to visualize the ring boundaries. Ring widths were measured to a precision of 0.01 mm using the TSAP measuring device (Rinntech). Tree-ring width (TRW) series were then visually cross-dated using standard dendrochronological methods (Stokes and Smiley, 1968) and checked for dating accuracy and measurement errors with the COFECHA program (Holmes, 1983). The tree-ring chronology was performed using the R-package ‘dplR’ (Bunn, 2008; R Core Team, 2022).

### 2.4 Triple isotope data

For stable isotope analysis, cores from five trees were selected, avoiding cores with visible defects. Measurements of δ^13^C, δ^18^O, δ²H were conducted separately for each selected tree and each year over the period 2000-2023. We split off the individual years with a razor blade under a stereomicroscope. For each year and tree, we pooled wood material from both cores per tree for a more representative tree value and to ensure sufficient material for measurement of small rings. The separated tree rings were transferred into filter bags (Ankom Technology, Macedon, NY, USA) for subsequent cellulose extraction. We performed cellulose extraction according to Boettger et al. (2007) and cellulose homogenization following Laumer et al. (2009). Cellulose samples were weighed (0.9-1.1 mg) into silver capsules, equilibrated with water vapor (Schuler et al., 2022), and then converted to H_2_ and CO by pyrolysis at 1420°C (PYRO-cube, Elementar, Hanau, Germany) for subsequent measurements of isotope ratios with an isotope-ratio mass spectrometer (IRMS; Delta Plus XP; Thermo Finnigan Mat, Bremen, Germany) with a precision ±0.2‰ for δ^13^C, ±0.3‰ for δ^18^O and δ^2^H 3.0‰ (Saurer et al., 2025).

In this study, δ^13^C was used as a proxy for intrinsic water-use efficiency and photosynthetic carbon assimilation (Farquhar et al., 1982; Scheidegger et al., 2000), δ^18^O as a proxy integrating source-water isotopic composition and evaporative leaf water enrichment (Roden et al., 2000; Barbour, 2007), and δ²H primarily as a proxy for source-water isotopic variability and physiological responses (Lehmann et al., 2021; Saurer et al., 2025).

### 2.5 _i_WUE inferred from carbon isotope discrimination

Measurements of δ^13^C were used to calculate iWUE for European beech trees. _i_WUE is defined as the ratio between the net photosynthesis rate (*A*) and the stomatal conductance to water vapor (g_s_) rate (Ehleringer & Dawson, 1992) and was computed as per Eqn 2:

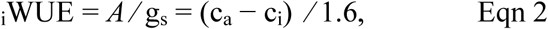

where c_a_ is the atmospheric CO_2_ concentration, c_i_ is the intercellular CO_2_ concentration (both in ppmv), and 1.6 represents the ratio of the diffusivities of water vapour and CO_2_ in air.

For European beech, _i_WUE can be derived from the carbon isotope composition (δ^13^C_t_ᵣₑₑ) of plant material, which is related to the isotopic composition of atmospheric CO_2_ (δ^13^Cₐ_tm_) and to the cᵢ/cₐ ratio, following the model by Farquhar and Richards (1984) and Farquhar et al. (1982) (Eqn 3):

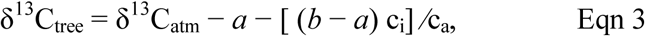

where *a* is the fractionation factor during diffusion through stomata (4.4‰) and *b* is the fractionation factor associated with Rubisco carboxylation (27‰). From Eqn 2, cᵢ can be derived as (Eqn 4):

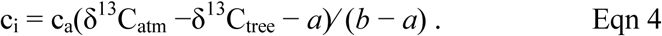

Finally, the _i_WUE is obtained as (Eqn 5):

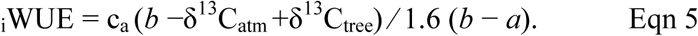

Values of δ^13^C_atm_ and the atmospheric concentration of CO_2_ for the period 2000–2023 were obtained from the Mauna Loa station data (available at http://www.esrl.noaa.gov/).

### 2.6 Modelling C and water cycle

Carbon and water fluxes were simulated using the 3D-CMCC-FEM (“*Three Dimensional - Coupled Model Carbon Cycle - Forest Ecosystem Module*”) (Collalti et al., 2016; 2020; 2024; Dalmonech et al., 2022; 2024; Puchi et al., 2026; Saponaro et al., 2025; 2026). The 3D-CMCC-FEM is a biogeochemical, biophysical and eco-physiological process-based model designed to simulate carbon and water cycle dynamics in forest ecosystems. The model has been extensively validated across species and climatic gradients within e.g., the Fluxnet, ICOS, and PROFOUND/ISIMIP networks (e.g., Marconi et al., 2017; Reyer et al., 2019, 2020; Mahnken et al., 2022).

Model simulations were used to estimate gross primary productivity (GPP), evapotranspiration (ET), intrinsic water-use efficiency (_i_WUE; the ratio of net carbon assimilation, *A*, to stomatal conductance for water vapour, *g_s_*), and ecosystem water-use efficiency (WUE_eco_; the ratio of GPP to ET). Simulated fluxes were compared with eddy-covariance observations and tree-ring isotope-derived estimates to evaluate the consistency of carbon–water coupling across daily, monthly and annual timescales.

Photosynthesis (*A*, µmol CO₂ m⁻² s⁻¹) is computed according to the biochemical model of Farquhar et al. (1980) with temperature acclimation as in Kattge & Knorr (2007) and soil moisture regulation as in Bonan et al. (2011). *A* is calculated as the minimum of two potential limitations (Eqn 6):

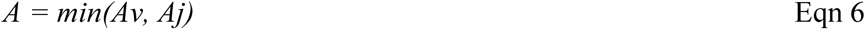

where Av is Rubisco-limited assimilation, and Aj is electron transport-limited assimilation,, computed respectively as (Eqn 7 and Eqn 8):

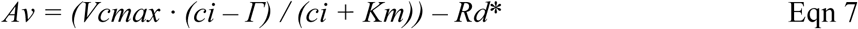

where *Vcmax* is the maximum carboxylation rate corrected for temperature and soil water availability, *ci* is intercellular CO_2_ concentration, *Γ* is the CO_2_ compensation point, *Km* is the Michaelis-Menten coefficient, and *Rd* is mitochondrial respiration in the light.

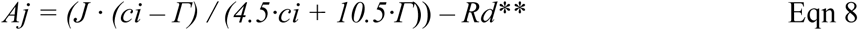

where *J* is the electron transport rate, and the empirical coefficients (4.5 and 10.5, respectively) follow Bernacchi et al. (2003).

Both *Vcmax* and *Jmax* (defined at 25°C) are dynamically adjusted using temperature response functions (Medlyn et al., 2002) and further downregulated by soil water content (Bonan et al., 2011). These parameters are computed separately for sunlit and shaded canopy fractions, incorporating nitrogen variation across canopy layers to ensure physiologically realistic limitations to photosynthesis through time.

Stomatal conductance (g_s_) is calculated daily following the multiplicative stomatal regulation model of Jarvis (1976) (Eqn 9):

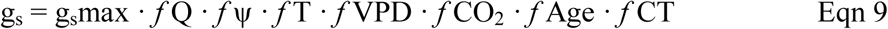

where *g_s_max* is the maximum potential stomatal conductance and *f* terms represent environmental and physiological modifiers (light, soil water potential, air temperature, vapor pressure deficit, atmospheric CO_2_ concentration, age, and temperature-pressure correction) varying from 0 (totally limitant) to 1 (not limitant), with the exception for *f*CO_2_, as described in Collalti et al. (2018; 2024) and Saponaro et al. (2026).

Intrinsic water-use efficiency is then derived from the ratio of *A* (Eqn 6) to *g*s (Eqn 9) as (Eqn 10):

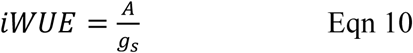

Daily model-derived _i_WUE^mod^ was averaged over the vegetative period and subsequently aggregated to annual values to ensure consistency with the annual integration of isotope-derived _i_WUE^obs^.

In the 3D-CMCC-FEM, gross primary productivity (GPP) is obtained by integrating canopy photosynthesis over time, while total evapotranspiration results from the combined simulation of canopy transpiration, soil evaporation, and interception/evaporation losses within the coupled water balance module. All evapotranspirative processes such as: canopy interception and evaporation (which includes also snow interception, melting and subsequent evaporation), stomatal/leaf/canopy transpiration, and soil evaporation, are computed at daily time steps using the Penman–Monteith equation as modified by Waring & Running (2007) and Monteith & Unsworth (2008). The formulation estimates latent heat fluxes as a function of incoming radiation, vapour pressure deficit, and surface conductances, based on meteorological conditions and surface characteristics (Collalti et al., 2024).

Soil moisture limitations directly constrain both photosynthetic assimilation and stomatal opening, thereby influencing _i_WUE dynamics, while climatic drivers including air temperature, vapour pressure deficit (VPD), and atmospheric CO_2_ concentration further modulate the trade-off between carbon gain and water loss through stomatal regulation (Puchi et al., 2026; Saponaro et al., 2026).

Model setup (Table 1), parameterization, and site-specific inputs (e.g., DBH-sapwood area relationships) are detailed in the Supplementary Material (Table S1). Simulations were performed using the R-package ‘R3DFEM’ (Collalti et al., 2022; Vangi et al., 2025).

**Table 1.**
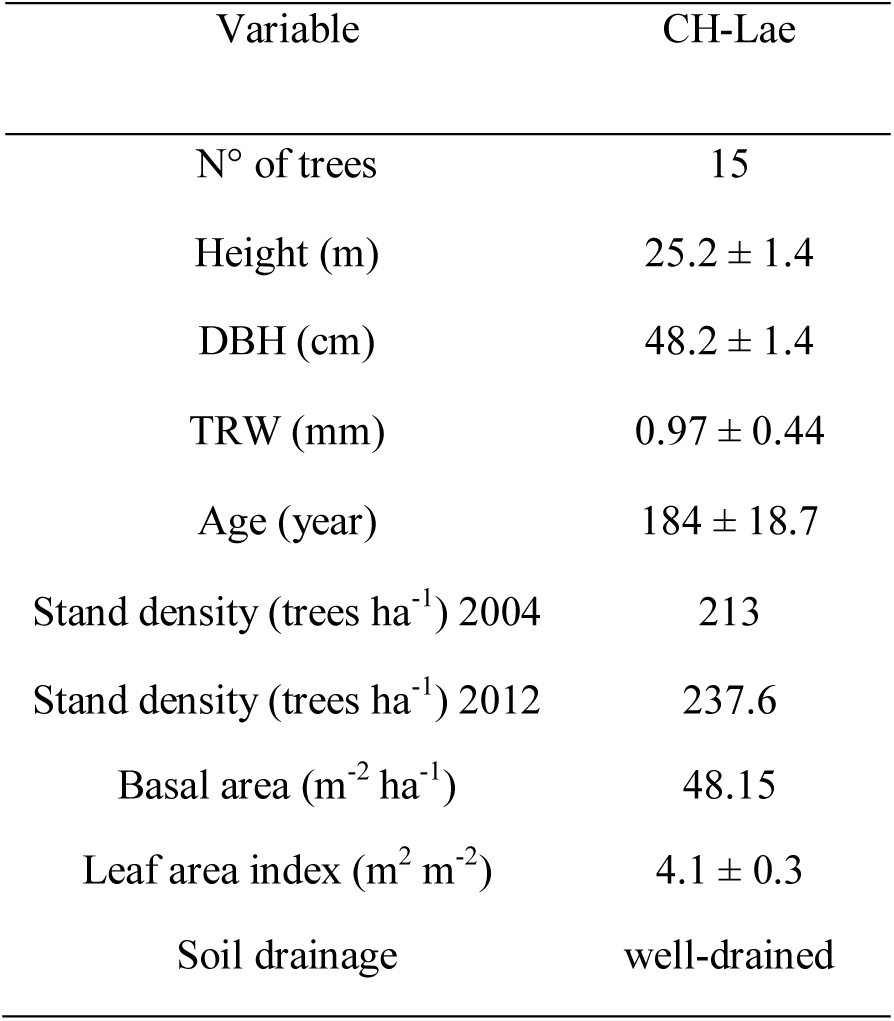
Biometric parameters of sampled *Fagus sylvatica* trees (N° of trees, height, DBH, TRW and age) and stand characteristics in 2004 for model initialization. Data are means ± standard deviation.

To validate the model, we first assessed the temporal relationships between modelled GPP and EC-derived GPP estimates across daily, seasonal, and interannual timescales (see Supplementary Notes S1; Figs. S1, S2, S3). This step confirmed that the simulated GPP dynamics reliably captured observed patterns, providing a robust basis for subsequent _i_WUE and WUE_eco_ analyses.

**Fig. 2.**
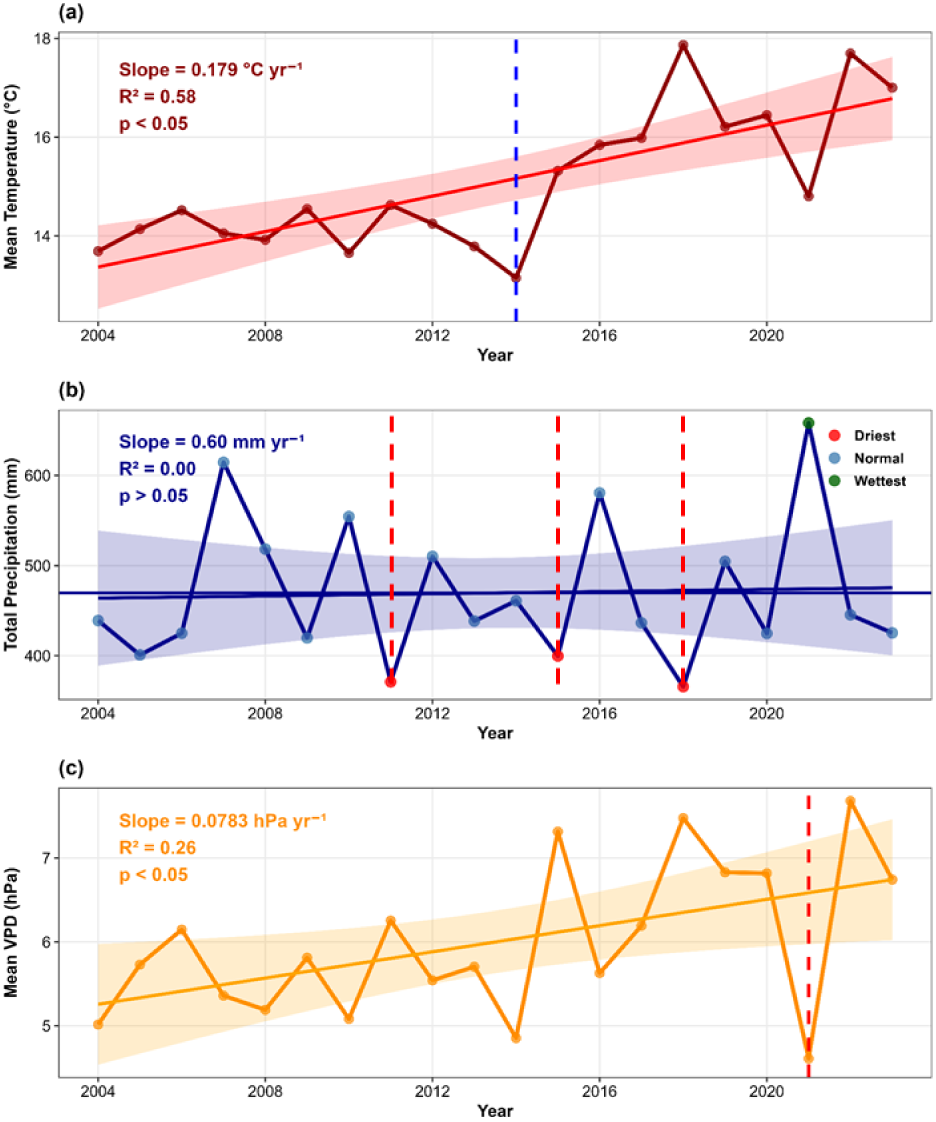
Climate trends during the growing season at the Lägeren CH-Lae flux tower data, Switzerland, for the period 2004-2023. (a) Mean annual temperature with linear regression trend line (red solid line) and 95% confidence interval (shaded area). The blue dashed vertical line indicates the coldest year (2014). (b) Total precipitation with linear regression trend line (dark blue solid line), mean precipitation reference line (horizontal dark blue line), and 95% confidence interval (shaded area). Data points are color-coded according to precipitation extremes: red circles indicate the three driest years, green circle indicates the wettest year (2021), and blue circles represent normal years. (c) Mean annual VPD with linear regression trend line (orange solid line) and 95% confidence interval (shaded area). The red dashed vertical line indicates the year with the lowest VPD (2021). Regression parameters (slope, R², *p*-value) are displayed at the top left for each variable.

**Fig. 3.**
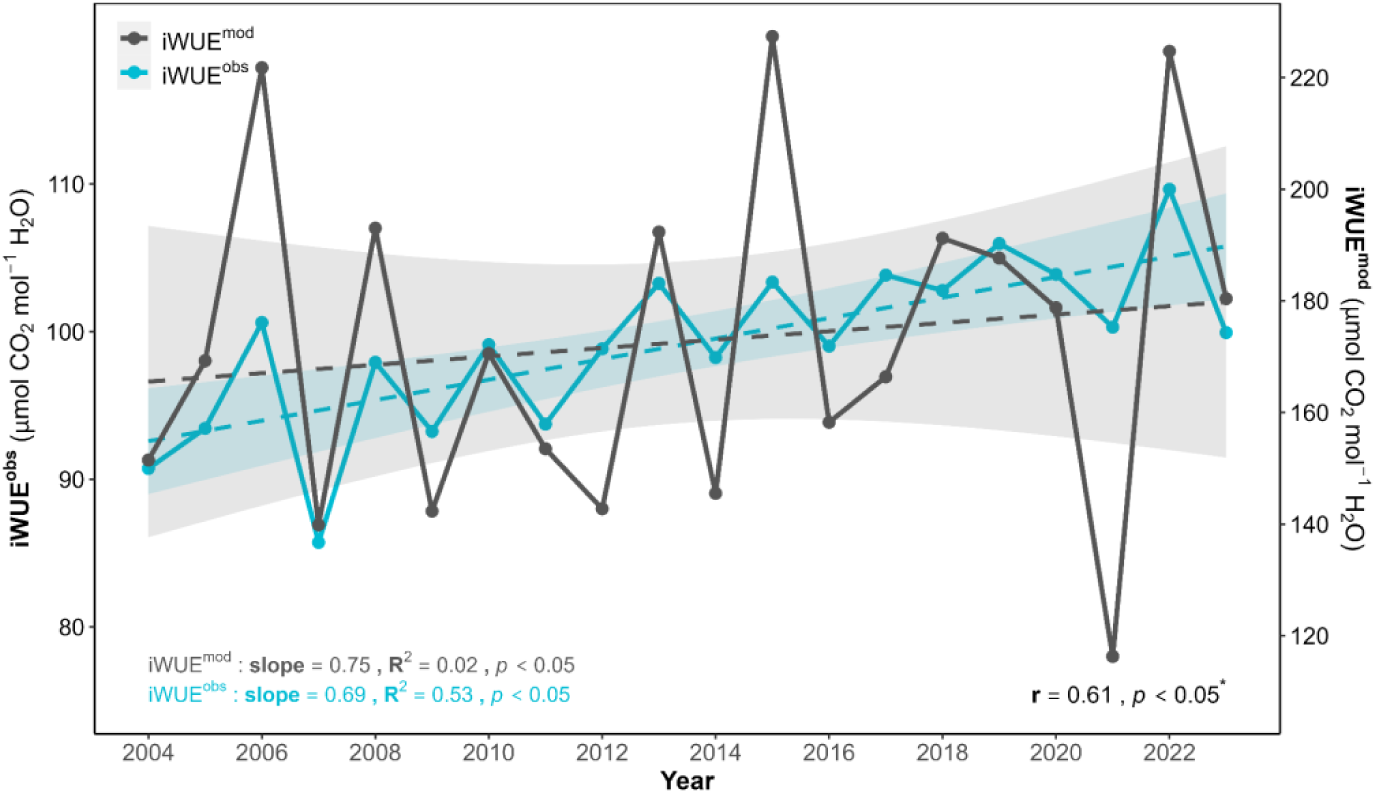
Inter-annual variability between _i_WUE^obs^ (cyan line) and _i_WUE^mod^ (gray line) from 21 June to 2 August (DOY 172-214) for the period 2004-2023. Dashed lines represent linear regression trends with shaded areas indicating 95% confidence intervals. Trend statistics (slope in µmol CO_2_ mol⁻¹ H₂O year⁻¹, coefficient of determination R², and *p*-value) are displayed adjacent to their corresponding trend lines in matching colors. Overall Pearson correlation coefficient and significance level between annual values are shown (bottom right, bold).

### 2.7. Statistical analyses and considerations

#### 2.7.1 Data cleaning, filtering, and computation of WUE_eco_ (ratio-of-sums)

Daily eddy-covariance and model outputs were quality-controlled prior to analysis. To refine the temporal scale of the analysis between _i_WUE derived from isotopic measurements and simulated _i_WUE, we applied a daily response function using the DendroTools R package (Jevšenak and Levanič, 2018). During the analysis, the observed and simulated _i_WUE time series were linearly detrended using the Standard Linear Detrending (SLD) method implemented in the *daily_response()* function to remove long-term trends and emphasize interannual variability. Pearson correlation coefficients were then calculated over moving windows ranging from 30 to 150 days within the growing season (DOY 105–243) to identify the period of strongest agreement between the two datasets. The highest positive correlation was observed between day of year (DOY) 172 (21 June) and DOY 214 (2 August) (Fig. S4). This interval largely overlaps with the June–August period previously identified as exhibiting the strongest climate–stable isotope relationships in *Fagus sylvatica* tree rings (Vitali et al., 2021). The identified period was subsequently used for comparisons between isotope-derived and modelled _i_WUE (Fig. S4).

To ensure internal consistency of model fluxes, days with GPP_model_ = 0 were forced to ET_model_ = 0. Extreme physiological values were removed using a daily percentile filter: for each date, the 99th percentile (P99) of _i_WUE and g_s_ was computed across all available records, and observations exceeding either threshold were discarded. Ecosystem water-use efficiency modelled (WUE_eco_^mod^) and obtained by eddy covariance observations (WUE_eco_^obs^) were derived from paired flux values with positive denominators, and daily values outside the range 0 < WUE_eco_ < 15 g C mm^-1^ H_2_O were excluded to prevent spurious spikes associated with very low ET.

Ecosystem water-use efficiency was computed as (Eqn 11):

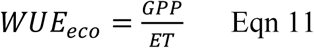

using a ratio-of-sums approach:

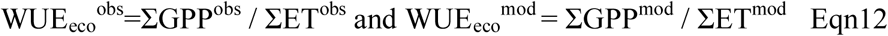

In this context ‘obs’ refers to eddy covariance measurements and ‘mod’ to model outputs, summing only over days with valid paired observations at daily, monthly, and annual scales. This approach avoids biases associated with averaging ratios of daily values (Ito & Inatomi, 2012) and enables cross-scale comparison between tree-level and ecosystem-level metrics. WUE_eco_ analyses were restricted to the growing season (15 April–31 August) for *Fagus sylvatica* at this site (Etzold et al. 2021). All statistical analyses were performed using R version 4.3.3 (R Core Team, 2024).

#### 2.7.2 Leaf-level water-use efficiency: isotope-derived vs. modelled *i*WUE

To evaluate the consistency between independent estimates of intrinsic water-use efficiency, isotope-derived _i_WUE^obs^ was compared with model-simulated _i_WUE^mod^ using Pearson correlation analysis and linear regression. Slope, coefficient of determination (R²), and 95% confidence intervals were computed for each regression model, and statistical significance was assessed using two-tailed *t*-tests (α = 0.05). Long-term temporal trends in both _i_WUE estimates were quantified by fitting linear regression models with *year* as the explanatory variable, with the regression slope representing the annual rate of change. The fitted trends were compared to assess the consistency between observed and modelled long-term trajectories over the study period (2004–2023).

#### 2.7.3 Ecosystem water-use efficiency: observed vs. modelled WUE_eco_

Agreement between eddy covariance-derived WUE_eco_^obs^ and model-simulated WUE_eco_^mod^ was evaluated using Pearson correlation analysis at multiple temporal scales (daily, monthly, and annual) to assess whether carbon–water coupling improved with increasing temporal aggregation. Linear regression models were applied to quantify the correspondence between observed and simulated ecosystem fluxes, with slope, R², and 95% confidence intervals reported for each scale. Days with missing paired observations were excluded prior to aggregation, following the ratio-of-sums approach described in Section 2.7.1.

#### 2.7.4 Isotopic controls on _i_WUE and radial growth

To interpret the physiological drivers of iWUE interannual variability, Pearson correlation and linear regression analyses were performed between _i_WUE^obs^ and the stable isotope ratios δ^18^O and δ²H measured in *F. sylvatica* tree-ring cellulose. These analyses were designed to distinguish between stomatal regulation and source water isotopic composition as the dominant controls on _i_WUE variability, following the conceptual framework of Scheidegger et al. (2000). According to this model, the covariation between δ^13^C and δ^18^O can be used to disentangle the relative contributions of stomatal conductance (*g*_s_) and mesophyll carboxylation capacity (*A*) to iWUE variability: when both δ^13^C and δ^18^O increase simultaneously, stomatal closure is the dominant driver, whereas when δ^13^C increases while δ^18^O remains unchanged or decreases, enhanced photosynthetic capacity (*A*) is the primary mechanism. To further characterise isotope coupling, the relationship between δ^13^C and δ^18^O was assessed to evaluate carbon–oxygen isotope co-variation, and the δ^18^O–δ²H relationship was examined to test for consistency with the meteoric water line and to evaluate the relative contribution of evaporative enrichment vs. source water dynamics to the oxygen and hydrogen isotope signals. Time series correlation between δ^18^O and simulated ecosystem stomatal conductance (*g*_s_) was additionally explored to identify physiological responses during climatically extreme years.

Finally, the relationship between TRW and isotope-derived _i_WUE was assessed using Pearson correlation and linear regression to evaluate whether long-term increases in water-use efficiency translated into enhanced radial growth. Additionally, the correlation between TRW and δ²H was examined across the full time series (2000-2023) to identify years in which source water availability or metabolic fractionation processes dominated the growth response, with particular attention to the extreme climatic conditions.

## 3. Results

### 3.1 Climate trends during the growing season

Mean growing season temperature increased significantly at 0.179 °C yr⁻¹ (R² = 0.58, p < 0.05; Fig. 2a), totaling ∼3.4 °C warming over the study period. The coldest growing season was 2014 (13.0 °C, blue dashed line), while the warmest years were 2018 (17.9 °C) and 2022 (17.6 °C). Instead, precipitation showed no significant trend (slope = 0.60 mm yr⁻¹, R² = 0.00, p > 0.05; Fig. 2b), fluctuating around a mean of 470 mm. The driest years were 2018 (363 mm), 2011 (369 mm), and 2015 (401 mm, red circles), while the wettest year was 2021 (665 mm, green circle), representing a 41% increase above the mean. Interestingly, mean vapor pressure deficit (VPD) increased significantly at 0.0783 hPa yr⁻¹ (R² = 0.26, p < 0.05; Fig. 2c), rising from ∼5.0 hPa (2004) to ∼6.8 hPa (2023), a 36% increase in atmospheric water demand. The lowest VPD occurred in 2021 (4.6 hPa, red dashed line), coinciding with the wettest growing season, while the highest VPD values were recorded in 2015 (7.2 hPa), 2018 (7.4 hPa), and 2022 (7.5 hPa). The combination of significant warming and increasing VPD without a corresponding precipitation trend indicates intensifying atmospheric water demand at the Lägeren site over the past two decades.

### 3.2 Intrinsic water-use efficiency: model-isotope comparison

Annual intrinsic water-use efficiency derived from tree-ring isotopes (_i_WUE^obs^) and simulated by the 3D-CMCC-FEM model (_i_WUE^mod^) were significantly correlated over the period 2004–2023 (r = 0.61, p < 0.05, n = 20; Fig. 3). The comparison was based on the optimal intra-annual window identified using DendroTools (DOY 172–214; 21 June–2 August; see Methods), indicating that model captures interannual variability in leaf-scale carbon-water coupling dynamics mediated by stomatal regulation (Fig. 3). _i_WUE^obs^ averaged 98.9 ± 6.2 µmol CO₂ mol⁻¹ H₂O, whereas _i_WUE^mod^ showed higher absolute values and substantially greater inter-annual variability (averaged 172.7 ± 30.3 µmol CO₂ mol⁻¹ H₂O). Both datasets exhibited significant positive trends over the study period, with _i_WUE^mod^ showing steeper increase (slope = 0.75 µmol CO₂ mol⁻¹ H₂O year⁻¹, R² = 0.02, *p* < 0.05) compared to _i_WUE^obs^ values (slope = 0.69 µmol CO₂ mol⁻¹ H₂O year⁻¹, R² = 0.53, *p* < 0.05).

### 3.3 Multi-temporal comparison of observed and modelled WUE_eco_

Daily WUE_eco_ during the growing season exhibited pronounced variability across 2004–2023 (Fig. 4). WUE_eco_^obs^ averaged 2.7 ± 1.4 g C mm⁻¹ H₂O, with episodic peaks to 10-15 g C mm⁻¹ H₂O during mid to late summer periods. WUE_eco_^mod^ showed higher mean 4.9 ± 2.7 g C mm⁻¹ H₂O. Despite this offset, daily values showed significant positive correlation (*r* = 0.26, *p* < 0.001, *n* = 2504), indicating that the model captured aspects of short-term meteorological forcing on carbon–water coupling.

**Fig. 4.**
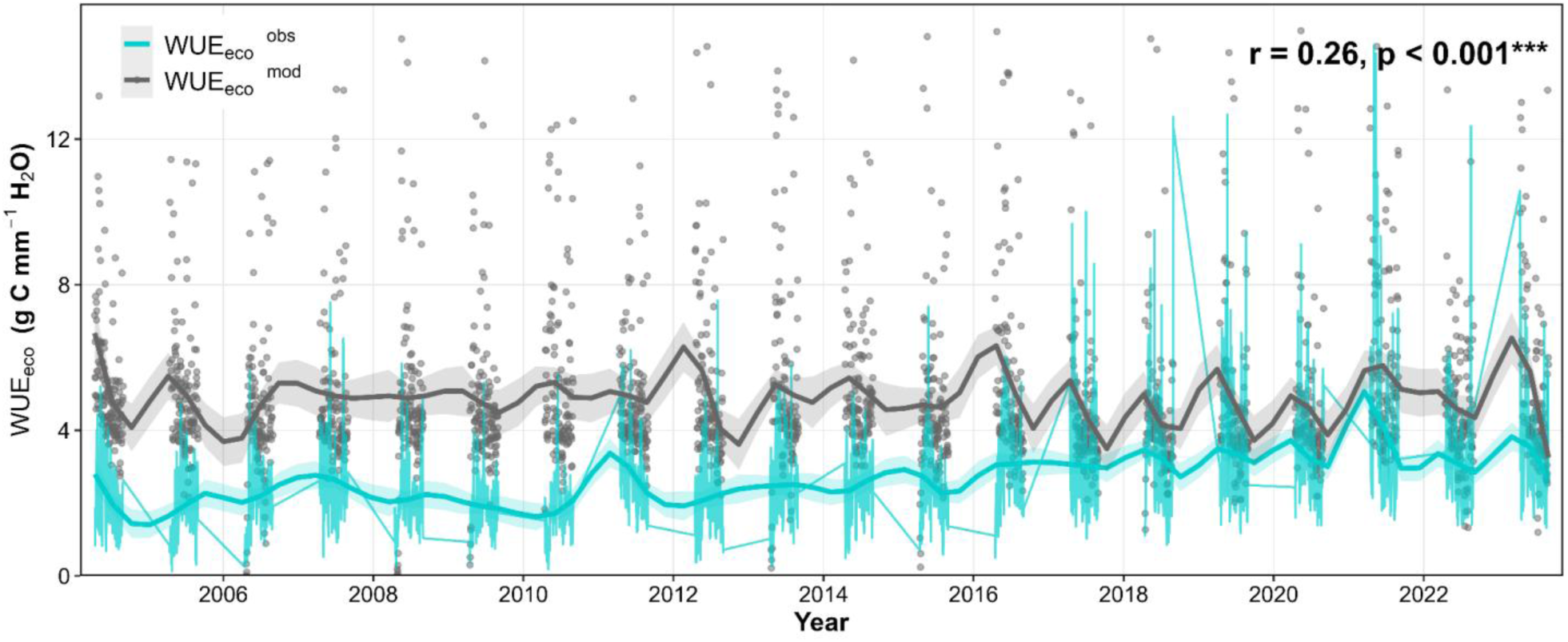
Daily comparison between WUE_eco_^obs^ (cyan lines) and WUE_eco_^mod^ (gray points) during the growing season in a mature *Fagus sylvatica* forest over two decades (2004-2023). Thick smooth lines represent LOESS trends (span = 0.1) highlighting temporal patterns. Data were filtered using the 99th percentile of growing-season WUE_eco_. The Pearson correlation coefficient (*r*) and significance level between observed and simulated values are shown (top right). *n* = 2504 daily observations.

LOESS-smoothed trajectories (span = 0.1) revealed distinct temporal patterns between datasets (Fig. 4). During 2004–2014, WUE_eco_^obs^ remained relatively low (∼2–3 g C mm⁻¹ H₂O) and stable, while WUE_eco_^mod^ fluctuated around a consistently higher mean (∼4–5 g C mm⁻¹ H₂O), with both series showing coherent synchronous seasonal oscillations throughout this period. From 2015 onwards, WUE_eco_^obs^ exhibited a progressive increase, gradually approaching the range of WUE_eco_^mod^, accompanied by intensified within-season variability and more frequent episodic peaks (10–13 g C mm⁻¹ H₂O) during mid-to-late summer. Both datasets captured concurrent responses to notable climatic events, including peaks during 2018 and 2021–2022.

At the monthly timescale, WUE_eco_ exhibited higher synchrony within the growing season (April-August) between WUE_eco_^obs^ and WUE ^mod^ across the 2004-2023 study period (Fig. 5). Observed monthly WUE_eco_^obs^ averaged 2.7 ± 0.9 g C mm⁻¹ H₂O, while WUE_eco_^mod^ outputs averaged 4.9 ± 1.2 g C mm⁻¹ H₂O, approximately double the observed values.

**Fig. 5.**
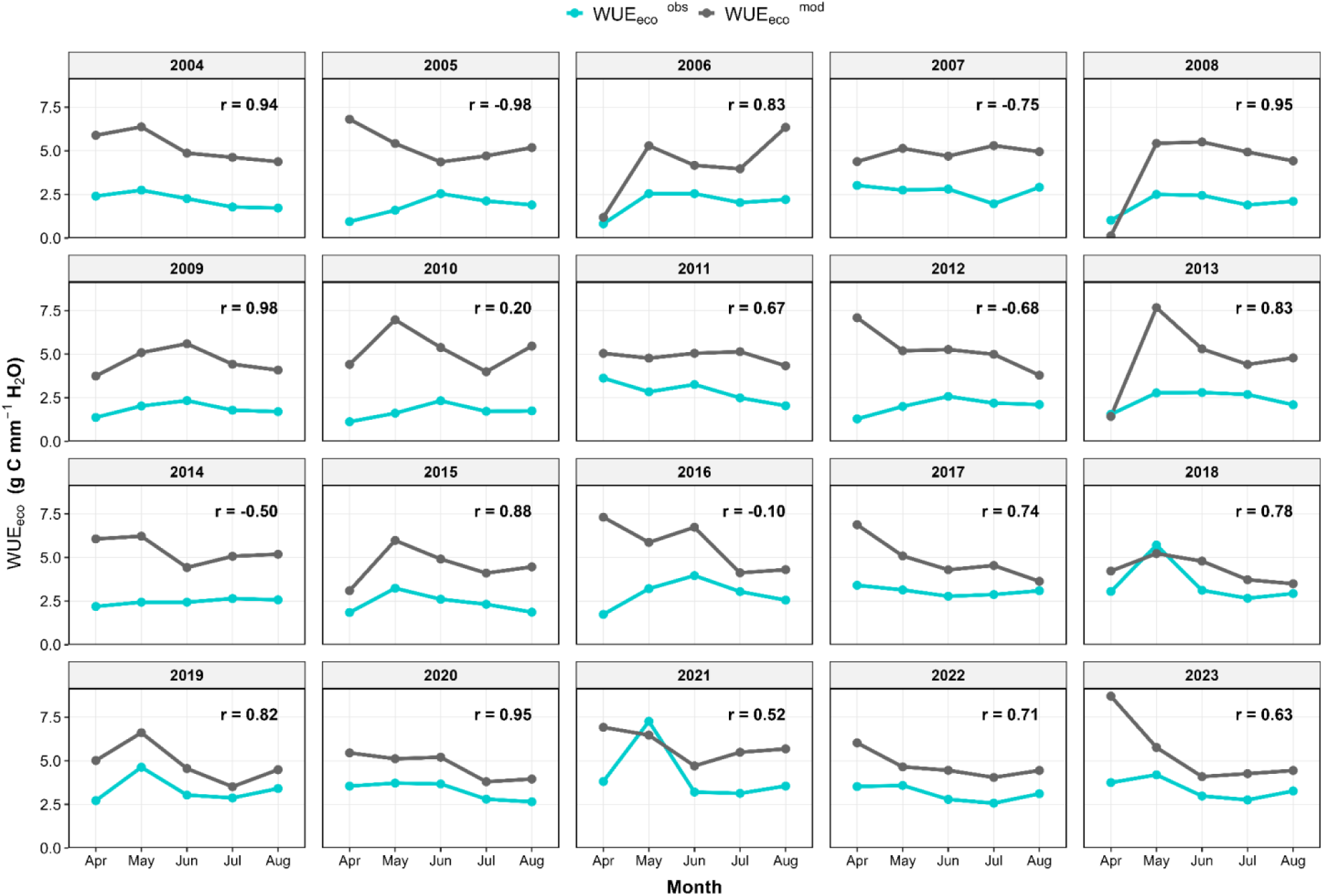
Monthly mean WUE_eco_ during the *F. sylvatica* growing season, 2004-2023, comparing WUE_eco_^obs^ (cyan line) with WUE_eco_^mod^ (gray line). Pearson correlation coefficients (*r*) are shown for each year.

Overall monthly correlation between WUE_eco_^obs^ and WUE_eco_^mod^ was statistically significant (*r* = 0.42, *p* < 0.001, *n* = 100 monthly means), reflecting high interannual variability in model–data agreement. Annual correlations ranged from *r* = −0.98 to *r* = 0.98, with 15 of 20 years showing positive correlations.

Temporal analysis revealed progressive improvement in model–data correspondence, with stronger agreement during 2015-2023 (*r* = 0.66, *p* < 0.001, *n* = 45) compared with 2004-2014 (*r* = 0.23, *p* = 0.003, *n* = 55). During years of strong positive correlation, both datasets captured concordant seasonal peaks during May–June and concordant minima during transitional months (April, August), consistent with the phenological constraints of the growing season at this temperate forest site.

Annual growing-season WUE^eco^ showed substantial differences between WUE_eco_^obs^ and WUE_eco_^mod^ (Fig. 6). The 3D-CMCC-FEM model overestimated WUE_eco_^mod^ (mean: 4.9 ± 0.3 g C mm⁻¹ H₂O) compared WUE_eco_^obs^ measurements (mean: 2.7 ± 0.6 g C mm⁻¹ H₂O). Temporal dynamics diverged markedly between datasets. WUE ^obs^ revealed a strong positive trend (slope = 0.083 g C mm⁻¹ H₂O year⁻¹, *p* < 0.001), with WUE_eco_^obs^ increasing ∼55% from 1.8 to 4.0 g C mm⁻¹ H₂O over the study period. Conversely, WUE_eco_^mod^ exhibited no significant trend (slope = −0.005, *p* = 0.716), fluctuating within a narrow range (4.3-5.7 g C mm⁻¹ H₂O) around a stable mean (Fig. 6).

**Fig. 6.**
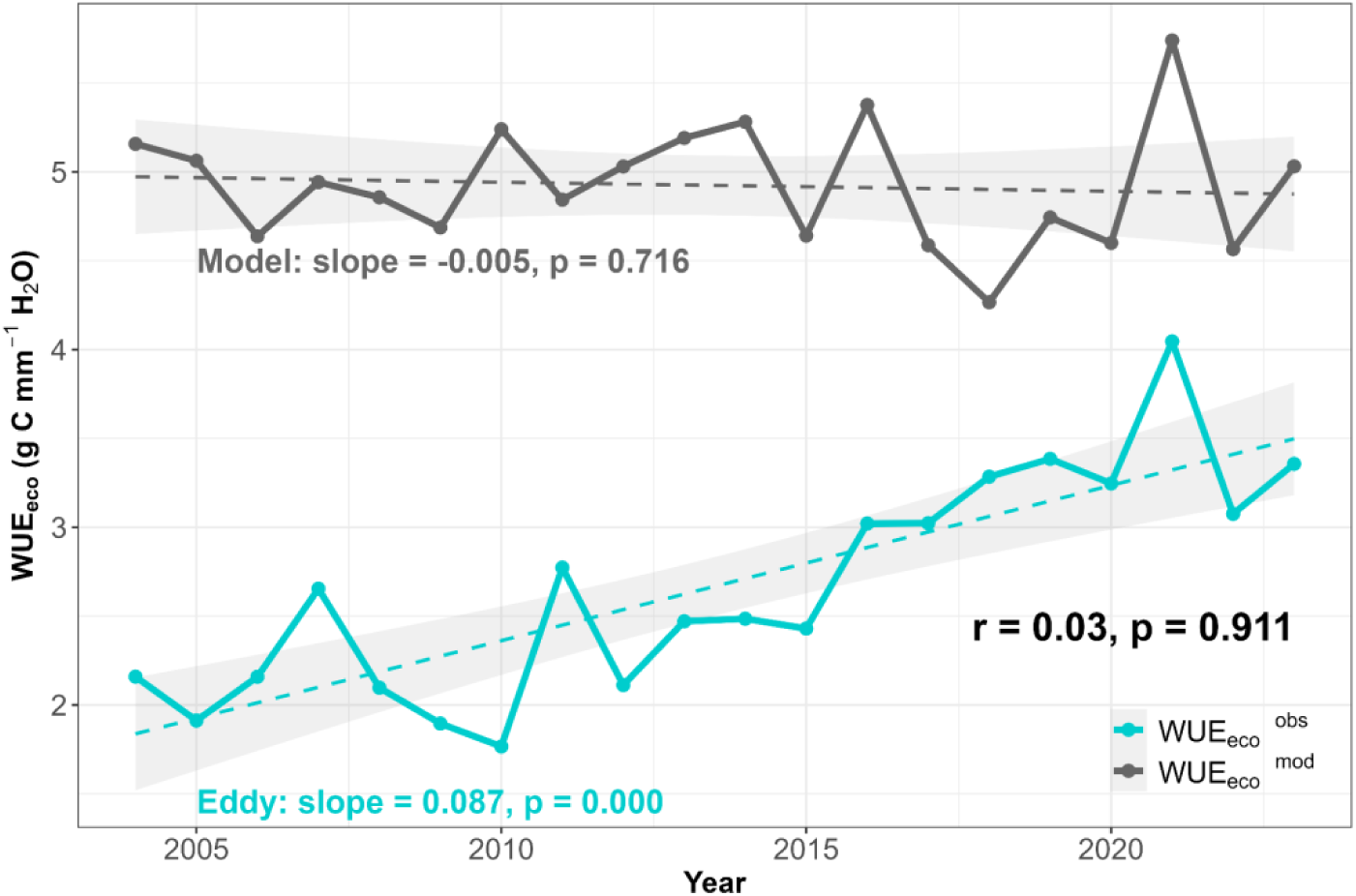
Annual WUE_eco_^obs^ from eddy covariance observations (cyan) and WUE_eco_^mod^ (gray) during the growing season of *F. sylvatica* for the period 2004-2023. Dashed lines indicate linear regression trends with shaded areas representing 95% confidence intervals. Trend statistics (slope and *p*-value) are displayed adjacent to their corresponding trend lines in matching colors. Bold text above the legend shows the Pearson correlation coefficient and significance level between observed and simulated annual values.

Although both time series displayed qualitatively similar interannual patterns from 2015 onwards—including synchronized minima (2017) and maxima (2021-2022), this partial convergence did not translate into overall agreement, with the correlation between annual values remaining weak and non-significant (*r* = 0.03, *p* = 0.911).

Interestingly, the comparison between annually resolved _i_WUE^obs^ and WUE_eco_^obs^ revealed a positive but non-significant relationship (R² = 0.22, slope = 0.045, p > 0.05; Fig. 7), indicating that despite their concurrent increasing tendencies over 2004-2023, the two metrics were not significantly coupled at the annual scale.

**Fig. 7.**
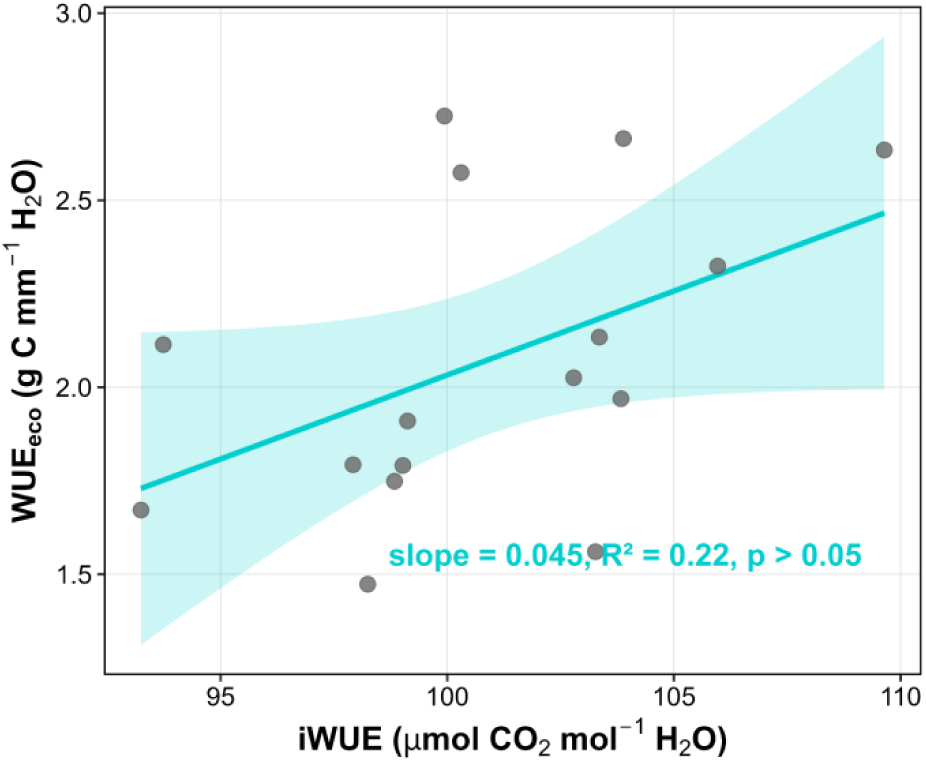
Relationship between annual _i_WUE^obs^ derived from tree-ring δ^13^C and annual WUE_eco_^obs^ derived from EC measurements at the Lägeren site (CH-Lae) over the period 2004-2023 (*n* = 20). The cyan line represents the linear regression fit with the shaded area indicating the 95% confidence interval. Regression statistics (slope, R², and *p*-value) are displayed within the panel.

### 3.2 Source water isotope controls on _i_WUE and TRW

The relationship between _i_WUE^obs^ and δ^18^O was negative but not statistically significant (R² = 0.15, slope = −4.132, p > 0.05; Fig. 8a). In contrast, _i_WUE^obs^ exhibited a stronger and significant negative relationship with δ²H (R² = 0.27, slope = −0.640, p < 0.05; Fig. 8b). Interestingly, δ²H and δ¹⁸ O in tree rings were significantly positive related (R² = 0.17, slope = 3.292, p < 0.05; Fig. 8c).

**Fig. 8.**
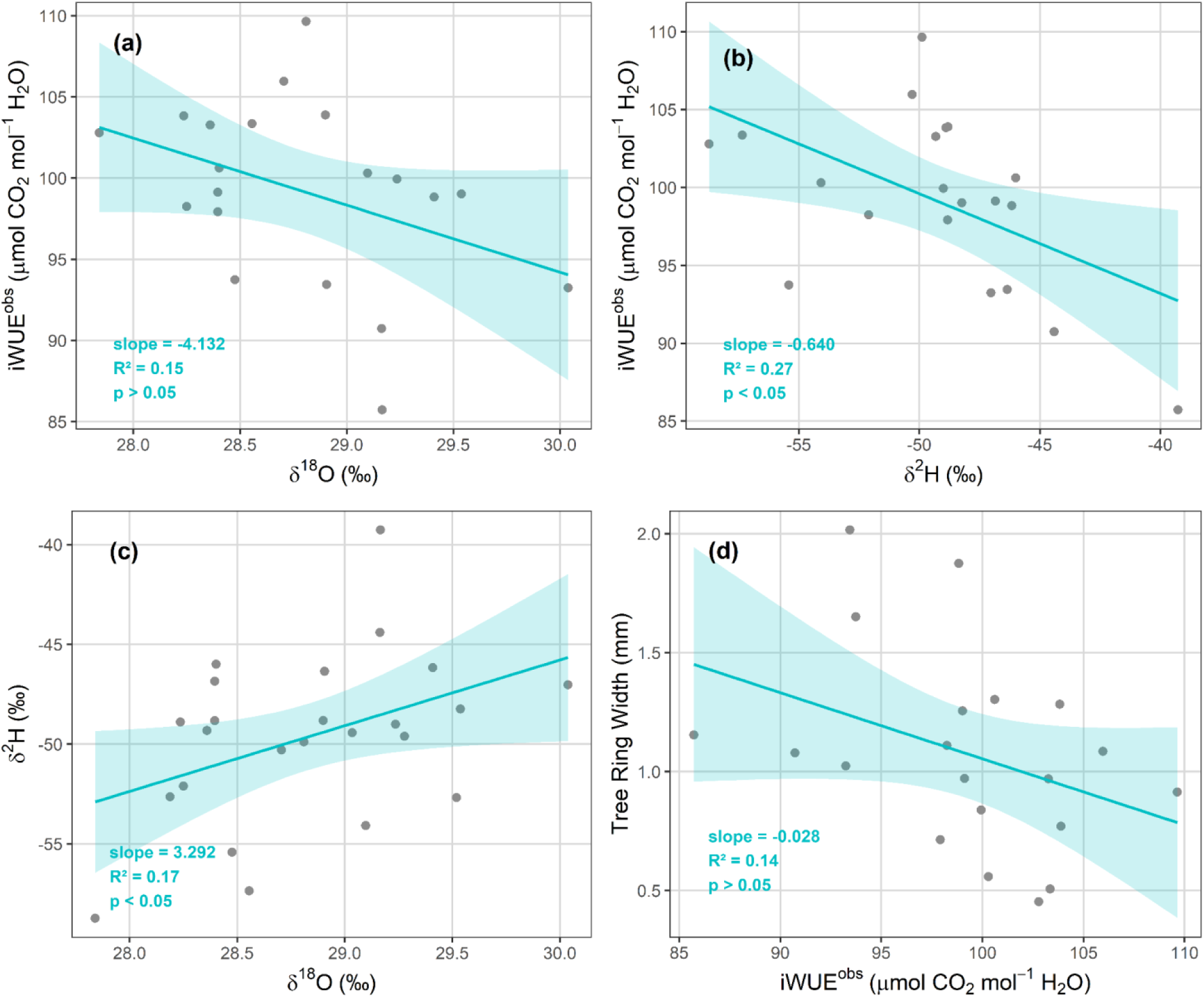
Relationships between intrinsic water use efficiency (_i_WUE^obs^) and isotopic composition of tree rings at CH-Lae for the period 2000-2023. (a) _i_WUE^obs^ vs. δ^18^O ‰); (b) _i_WUE^obs^ vs. δ²H (‰); (c) δ²H vs. δ^18^O; (d) TRW vs _i_WUE^obs^. Linear regression lines with 95% confidence intervals (shaded areas in cyan) are shown. Regression parameters (slope, R², *p*-value) are displayed at bottom left in cyan. Each point represents an individual year (*n* = 24).

TRW showed a weak negative but non-significant relationship with _i_WUE^obs^ (R² = 0.14, slope = −0.028, p > 0.05; Fig. 8d).

Additional insight from triple-isotope relationships revealed a decoupling between carbon and oxygen isotope signals, as indicated by the weak and non-significant relationship between δ^13^C and δ^18^O (R² = 0.02, p > 0.05; Fig. S5a). In contrast, δ^13^C exhibited a stronger and significant positive relationship with δ²H (R² = 0.29, p < 0.01; Fig. S5b).

Interestingly, TRW correlated positively and significantly with δ²H (r = 0.42, *p* < 0.05; Fig. 9a), most noticeable during the extreme drought year of 2011 and the coldest year 2014 (Fig. 9a). In contrast, TRW did not correlate significantly with either δ^18^O (r = 0.15, *p* > 0.05; Fig. 9b) or δ^13^C (r = 0.39, *p* > 0.05; Fig. 9c).

**Fig. 9.**
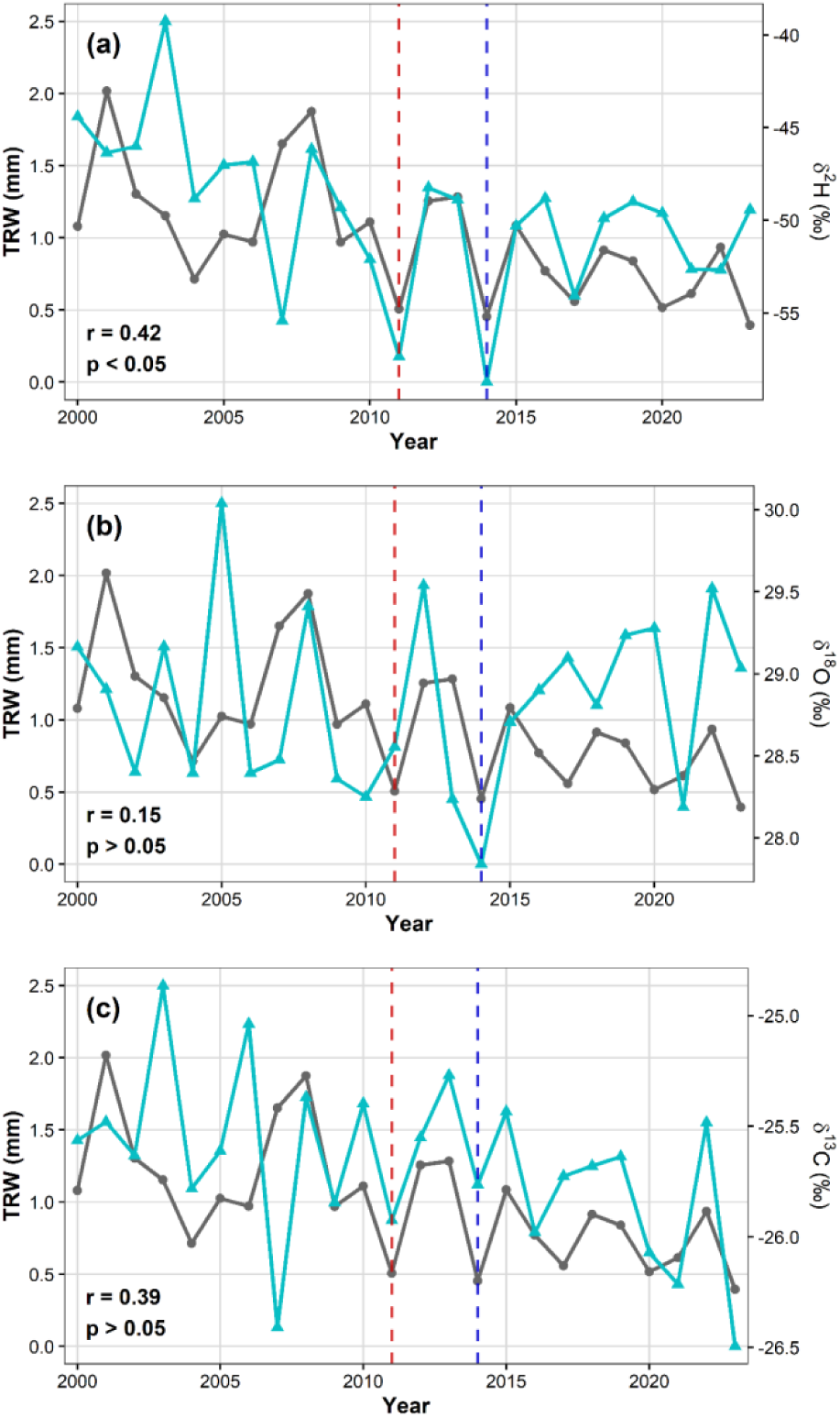
Interannual time series of tree-ring width (TRW, gray lines; right axis) and stable isotopes ratios (cyan lines; right axis) of *F. sylvatica* from 2000-2023: (a) δ²H, (b) δ^18^O and (c) δ^13^C. The vertical red dashed lines indicate the driest years on record (2011) and the vertical blue dashed line the coldest (2014), both identified from meteorological data at the CH-Lae flux tower.

Temporal analysis of δ^18^O and simulated ecosystem stomatal conductance (*g*_s_) showed that the lowest δ^18^O values coincided with the highest simulated *g_s_* during the exceptionally wet summer of 2021 (476 mm precipitation; Fig. 2 and Fig. S6). During this year, δ^18^O declined to approximately 28.2‰ while *g_s_* reached its maximum (∼0.038 mol m⁻² s⁻¹), consistent with the expected physiological response to abundant water availability.

## Discussion

### 4.1 Consistency between modelled and observed _i_WUE

Our multi-proxy analysis revealed a consistent increase in _i_WUE over the last two decades derived both from tree-ring δ^13^C and simulations of the process-based model 3D-CMCC-FEM. The positive _i_WUE trend is consistent with the widely documented forest response to rising atmospheric CO_2_ across temperate European beech forests, with stronger responses reported in water-limited sites compared to mesic environments (Frank et al., 2015; Guerrieri et al., 2019; Wang et al., 2025). Our findings show an approximately 20% increase in _i_WUE over two decades at the Lägeren site, consistent with these regional patterns (Gharun et al., 2021). The concurrent significant warming and increasing VPD documented at the site, without a corresponding trend in precipitation, suggest that the _i_WUE increase reflects both CO_2_ fertilization and physiological responses to increasing atmospheric water demand (Puchi et al., 2024; Scapucci et al., 2025).

The significant positive correlation between isotope-derived and modelled _i_WUE indicates that both approaches consistently captured the interannual dynamics of _i_WUE (Saponaro et al., 2025; 2026). However, rather than validating one another, they should be regarded as complementary and independent estimates of leaf-level carbon–water coupling, each integrating physiological processes over different temporal scales and with different methodological assumptions. The agreement in temporal trends therefore provides converging evidence that iWUE has increased over the last two decades in this temperate beech forest, while differences in absolute values reflect the inherent characteristics of each approach rather than inconsistencies in the underlying physiological response.

Observed _i_WUE values ranged between 80 and 110 μmol mol⁻¹, consistent with reported values for temperate forest species under summer conditions (Medlyn et al., 2011; Keenan et al., 2013). In contrast, modelled _i_WUE averaged approximately 172 µmol CO₂ mol⁻¹ H₂O and exhibited substantially greater interannual variability. This systematic offset in magnitude is unlikely to reflect a contradiction between the two approaches, but rather differences in the temporal scales over which _i_WUE is represented. Whereas isotope-derived _i_WUE integrates physiological responses throughout the wood-formation period, modelled _i_WUE was calculated from daily simulations within the optimal intra-annual window identified using DendroTools (21 June–2 August), corresponding to the period of maximum summer temperatures and atmospheric evaporative demand. During this period, elevated temperature and VPD simultaneously enhance photosynthetic assimilation while reducing stomatal conductance, leading to considerably higher instantaneous _i_WUE values than those preserved in annually integrated tree-ring isotope records (Medlyn et al., 2017; Grossiord et al., 2020; Novick et al., 2016). Consequently, daily model simulations during peak summer conditions are expected to produce substantially higher _i_WUE values than annually integrated tree-ring isotope records, which reflect the cumulative physiological response over the wood-formation period.

Therefore, comparisons between isotope-derived and modelled _i_WUE should primarily emphasize temporal variability and long-term trends rather than direct agreement in absolute values. Despite the systematic offset, both approaches consistently identified increasing _i_WUE through time, suggesting that they capture common physiological responses to rising atmospheric CO_2_ and increasing atmospheric water demand while integrating these responses over different temporal scales. Similar discrepancies have been reported in previous studies comparing isotope-derived physiological proxies with process-based model simulations, highlighting the importance of accounting for temporal aggregation and methodological differences when interpreting _i_WUE (De Kauwe et al., 2013; Vitali et al., 2025).

### 4.2 Decoupling between leaf-level and ecosystem-level water-use efficiency

Despite the strong agreement observed at the leaf scale, our results show that WUE_eco_ derived from EC exhibits markedly different magnitudes and temporal variability compared with _i_WUE inferred from tree rings. This mismatch highlights the fundamental scale dependency of WUE metrics, which integrate different processes and spatial domains, temporal scales, and critically species compositions (Li et al., 2022; Grossiord et al., 2020). Whereas _i_WUE reflects leaf-level physiological regulation integrated during wood formation, WUE_eco_ represents an emergent property of vegetation–atmosphere interactions across the entire canopy (Medlyn et al., 2017). Critically, the EC signal represents a bulk integrated flux from the entire tower footprint, encompassing all species, soil evaporation, understory vegetation, and canopy interception, making direct comparison with a single-species model inherently challenging (Knauer et al., 2018; Baldocchi, 2003). Consequently, differences between _i_WUE and WUE_eco_ should primarily be interpreted in terms of the processes and temporal scales represented by each metric rather than as inconsistencies between the two approaches. Flux-based WUE_eco_ therefore reflects both physiological controls on carbon assimilation and all hydrological processes operating at canopy and soil scales, which may decouple ecosystem-level signals from leaf-level stomatal regulation (Beer et al., 2009; Keenan et al., 2013; Paul-Limoges et al., 2017; Puchi et al., 2025).

The divergence between WUE_eco_^obs^ and WUE_eco_^mod^ is primarily attributable to a systematic underestimation of ET by the model (approximately 50% of observed values), rather than to differences in GPP, which remained broadly comparable between the two approaches (Fig. S1). By excluding the water use of co-occurring broadleaf species, evergreen conifers, younger trees, and understory vegetation, the model omits key contributors to canopy transpiration, interception losses, and soil water uptake, leading to systematically inflated WUE_eco_^mod^ values (Anderegg et al., 2019; Grossiord et al., 2020). Above-canopy EC measurements at Lägeren may additionally be influenced by advection effects associated with the site’s complex topography, and standard friction velocity-filtering procedures were applied to minimize these effects (Etzold et al., 2010; Aubinet et al., 2010). An additional and likely source of discrepancy between WUE_eco_^obs^ and WUE_eco_^mod^ arises from differences in species representativeness. The model was parametrized to simulate an old-growth deciduous stand composed exclusively of mature *F. sylvatica*, whereas EC measurements integrate fluxes across the entire mixed-species stand, including *F. excelsior* (∼20% basal area), *Acer* sp. (∼20%), *A. alba* (∼10%), and understory vegetation dominated by *Allium ursinum* in spring and early summer. The full quantification of these non-transpirational and multi-species water fluxes is addressed in Section 4.3.

The divergent temporal trends, with observations showing a significant positive trend in WUE_eco_^obs^ (slope = 0.087, p < 0.001) while model simulations remained stable (slope = −0.005, p = 0.716), further support this interpretation. The observed trend likely reflects changes in ecosystem water use dynamics at CH-Lae, driven by the documented increase in VPD (+36% over the study period) and differential canopy-level physiological responses across species. In this context, the coupling between *A* and *g*_s_ is not static but varies with temperature and VPD, and incorporating thermal sensitivity of stomatal conductance provides important improvements to models of plant carbon-water exchange under future climate conditions (Chessman et al., 2026). Moreover, increasing atmospheric demand can partially decouple T from *g_s_*, as T may continue to increase with rising VPD despite progressive stomatal closure because of the steeper leaf-to-air vapor pressure gradient (Diao et al., 2024). This nonlinear response implies that ecosystem water loss does not necessarily scale proportionally with stomatal regulation, thereby contributing to divergence between leaf-level _i_WUE and ecosystem-scale WUE_eco_ under high VPD conditions. The presence of evergreen species, younger trees, and a seasonally dense understory during spring and early summer may have further amplified these differential responses, as these functional groups exhibit distinct stomatal regulation strategies compared to mature deciduous *F. sylvatica* (Anderegg et al., 2019; Scapucci et al., 2025; Shekhar et al., 2025). Since the model simulates only the deciduous old-growth component using a static stomatal conductance parameterization, it cannot capture these thermally-driven and species-specific responses, explaining its inability to reproduce the observed positive trend in WUE_eco_^obs^.

Further, hydrological partitioning of ET may represent an additional constraint on comparing physiological and ecosystem-scale WUE metrics (Paul-Limoges et al., 2020). While T is directly controlled by stomatal conductance and therefore linked to photosynthesis, ET measured by EC also includes soil evaporation and canopy interception (Knauer et al., 2018; Nelson et al., 2022) and forest management (Saponaro et al., 2026).

T represents between 40% and 80% of total ET in temperate forests, depending on canopy structure, leaf area index, soil moisture availability, and atmospheric demand (e.g., Jasechko et al., 2013; Paul-Limoges et al., 2020). At forest sites with dense canopies and frequent precipitation, such as Lägeren, interception losses can account for 15-30% of total ET, particularly during the full growing season. Furthermore, based on unpublished below-canopy eddy covariance measurements at Lägeren (L. Hörtnagl, pers. comm.), understory ET contributes approximately 10% of the above-canopy flux, a component entirely absent from the single-species model simulation. Taken together, these non-transpirational and understory fluxes represent a substantial and systematic source of ET underestimation in the model, directly explaining the magnitude offset in WUE_eco_ documented in Section 4.2.

Because these water fluxes are not coupled to photosynthetic carbon assimilation by *F. sylvatica*, WUE_eco_ calculated as GPP/ET is expected to be systematically lower than plant-level metrics derived from stomatal processes alone (Knauer et al., 2018; Gong et al., 2022). This mathematical distinction, combined with the structural limitation of the single-species model configuration, provides a comprehensive explanation for the systematic WUE_eco_ offset observed in our study, without invoking differences in carbon assimilation between approaches.

Improving ET partitioning in process-based models therefore represents a critical step toward reconciling leaf-level and ecosystem-scale WUE estimates. Future model developments should incorporate multi-species canopy representations, explicit treatment of evergreen and early-successional components, understory vegetation modules, and improved interception and soil evaporation schemes to more realistically simulate the full water balance of structurally complex mixed-species temperate forests (Knauer et al., 2018; Nelson et al., 2022; Saponaro et al., 2026).

Similar discrepancies between leaf-level and ecosystem-level WUE have been reported in previous studies combining tree-ring isotopes and EC observations (Dekker et al., 2016; Lavergne et al., 2019; Li et al., 2022), but the role of species composition, understory contributions, and ET partitioning in driving these mismatches has only recently been recognized as a central challenge (Wehr et al., 2016; Medlyn et al., 2017).

### 4.3 Temporal scale-dependent dynamics of ecosystem WUE

The comparison between EC and modelled WUE_eco_ further revealed that model-data agreement improved with temporal aggregation. Daily WUE_eco_ exhibited weak correlations, reflecting the strong sensitivity of short-term fluxes to rapid fluctuations in solar radiation, VPD, and atmospheric turbulence. High-frequency meteorological variability can strongly influence stomatal conductance and canopy transpiration, generating short-term fluctuations that are difficult for ecosystem models to reproduce accurately (Knauer et al., 2018; Grossiord et al., 2020). Additionally, daily EC measurements at Lägeren are subject to uncertainty arising from gap-filling procedures, energy balance closure issues, and advection effects associated with the site’s complex topography. Advection has been estimated to account for up to 36% of net ecosystem productivity at this site, and standard u-filtering was applied to minimize these effects, although residual advective losses may still influence daily flux estimates (Baldocchi, 2003; Etzold et al., 2010; Aubinet et al., 2012; Medlyn et al., 2017).

These results are consistent with recent multi-scale studies showing that WUE depends on the biological scale at which it is evaluated. Rambal et al. (2025) similarly reported contrasting drought responses between _i_WUE and WUE_eco_. Our results extend this perspective by integrating tree-ring isotopes, eddy covariance observations, and process-based modelling to provide a mechanistic interpretation of these scale-dependent differences.

When aggregated to monthly timescales, the agreement between observed and simulated WUE_eco_ improved substantially, indicating that seasonal patterns of carbon-water coupling are more reliably reproduced by the model than short-term variability (Ito & Inatomi 2012). This improvement reflects the fact that monthly aggregation averages out high-frequency meteorological noise and measurement errors, allowing the underlying physiological responses to temperature, radiation, and soil moisture cycles to emerge more clearly (Medlyn et al., 2017; Lavergne et al., 2019).

At this temporal scale, the model’s representation of phenological development, seasonal acclimation of photosynthetic capacity, and progressive soil water depletion appears adequate to capture the dominant controls on ecosystem carbon-water coupling. The synchronous responses to climatic events, such as the 2018 drought (Gharun et al., 2020) and the 2021 wet year, further demonstrate that the model reproduces key aspects of interannual variability when integrated over monthly periods (Reyer et al., 2020; Collalti et al., 2016).

### 4.4 Disentangling physiological and hydrological signals driving WUE trends across scales

The triple isotope analysis (δ^13^C, δ^18^O, δ²H) of tree rings provided complementary insights into the mechanisms underlying long-term changes in _i_WUE and WUE_eco_ trends at the Lägeren site. Rather than reflecting a single physiological process, the three isotopes integrate different components of the soil–plant–atmosphere continuum. Consistent with previous studies (Scheidegger et al., 2000; Gessler et al., 2014; Vitali et al., 2022), our results suggest that carbon and water-related isotope signals become increasingly decoupled under mesic conditions, emphasizing that long-term _i_WUE trends cannot be interpreted solely in terms of stomatal regulation but must also account for hydrological controls associated with source-water uptake and transport.

The isotope evidence provides important mechanistic insights into the processes underlying the observed increase in _i_WUE (Gessler et al., 2014; Saurer & Voelker, 2022). If the long-term increase had been driven primarily by progressive stomatal closure, increasing evaporative enrichment of leaf water would be expected to produce higher δ^18^O values in tree-ring cellulose (Scheidegger et al., 2000; Barbour et al., 2004). However, neither δ^18^O nor δ²H exhibited patterns consistent with progressive isotopic enrichment, indicating that sustained reductions in stomatal conductance were unlikely to be the dominant mechanism underlying the observed increase in _i_WUE. Instead, our results suggest that the increase in _i_WUE was driven predominantly by enhanced photosynthetic carbon assimilation (A), or by a combination of increased assimilation and relatively modest adjustments in stomatal conductance, consistent with the conceptual dual-isotope framework proposed by Scheidegger et al. (2000).

The positive and significant relationship between δ²H and δ^18^O in tree rings of *F. sylvatica* (R2 = 0.27) is stronger than reported from longer European chronologies (Vitali et al., 2022), suggesting that both isotopes were governed by similar hydrological processes at the LAE site. Moreover, this coupling indicates that trees remained well acclimatized to the prevailing environmental conditions during the study period (Vitali et al., 2024). Although δ²H is generally considered less responsive than δ^18^O to evaporative enrichment associated with atmospheric demand it can also be influenced by post-photosynthetic metabolic fractionation under drought or physiological stress, leading to a decoupling between δ²H and δ^18^O signals (Cormier et al., 2018; Lehmann et al., 2021). However, the significant positive relationship observed in our study provides little evidence that such metabolic processes dominated δ²H variability during the study period. Instead, the coupled behaviour of δ²H and δ^18^O suggests that both isotopes were primarily controlled by hydrological processes, including source-water isotopic composition and leaf-water enrichment during tree-ring formation. This interpretation is consistent with Brinkmann et al. (2018), who demonstrated that xylem water integrates both recent precipitation and older soil water stored over several months, with winter precipitation contributing substantially to transpiration during the subsequent growing season in *F. sylvatica* at CH-Lae. Consequently, interannual variability in precipitation inputs, soil water residence time, and rooting depth can exert a dominant control on the isotopic composition of source water before isotopic signals are modified by leaf-level physiological processes.

Consequently, the negative relationships between δ^13^C-derived _i_WUE and both δ^18^O and δ²H suggest that leaf physiological processes are closely linked to source-water origin and uptake dynamics (e.g., recent precipitation versus shallow or deeper soil water) at the CH-Lae site. During relatively dry conditions, enhanced reliance on deeper soil water reserves, which are generally more isotopically depleted because they integrate older precipitation inputs, may simultaneously increase _i_WUE while producing lower δ^18^O and δ²H values in tree rings (Brinkmann et al., 2018). These findings indicate that interannual variability in source-water isotopic composition largely overprints physiological signals in both isotopes under the mesic conditions prevailing at Lägeren.

Nevertheless, this dominance of hydrological controls does not exclude a physiological contribution to δ²H. Unlike δ^18^O, hydrogen isotopes are increasingly recognized as integrating both hydrological processes and post-photosynthetic metabolism, including carbohydrate remobilization and carbon allocation (Lehmann et al., 2021; Vitali et al., 2022). Their Europe-wide synthesis demonstrated that δ²H is a weaker climatic proxy than δ^13^C or δ^18^O because it contains a substantial physiological component related to metabolic fractionation. Therefore, although source-water variability appears to dominate δ²H at the CH-Lae site under average conditions, physiological processes may become increasingly important during years characterized by exceptional drought or other environmental stress, when changes in carbon allocation and the utilization of stored assimilates are expected to contribute more strongly to the hydrogen isotope signal. This mechanism could explain deviations from the otherwise strong hydrological coupling between δ²H and δ^18^O observed during extreme years (Holloway-Phillips et al., 2026).

Moreover, assuming that δ^18^O in tree rings primarily reflects source-water isotopic composition rather than evaporative enrichment, our results suggest a relatively low sensitivity of δ^18^O to leaf-water enrichment and stomatal regulation at CH-Lae. This interpretation is supported by the negative relationship between δ^18^O and model-simulated *g_s,_* with appreciable sensitivity observed only during the exceptionally wet year of 2021. Likewise, the three driest growing seasons (2011, 2015 and 2018) produced only modest changes in both δ^13^C and δ^18^O (Fig. 9), indicating that severe summer drought did not induce the pronounced isotopic enrichment expected under sustained stomatal limitation. This muted response is consistent with the interpretation that mature *Fagus sylvatica* buffered drought through continued access to relatively stable soil water sources, thereby maintaining leaf water status and reducing evaporative enrichment despite high atmospheric demand Consequently, the weak relationship between δ^13^C and δ^18^O limits the direct application of the dual-isotope conceptual framework for separating the relative contributions of photosynthetic assimilation (A) and stomatal conductance (*g_s_*) at this site. (Scheidegger et al., 2000; Siegwolf et al., 2023). Instead, the limited sensitivity of δ^18^O to *g_s_* suggests that the observed increase in _i_WUE over the last two decades was driven primarily by enhanced photosynthetic carbon assimilation rather than sustained stomatal closure. This interpretation is directly supported by the significant positive trend in GPP observed over the study period, in both eddy covariance (slope = 18.98, *p* < 0.001) and model estimates (slope = 24.20, *p* < 0.001), with strong correlation between the two approaches (*r* = 0.63, *p* < 0.01; Fig. S1). The concurrent increase in GPP and _i_WUE suggests that enhanced CO_2_ fertilization sustained carbon assimilation capacity, while the isotopic composition of source water, rather than stomatal regulation, remained the dominant control on δ^18^O and δ²H variability at this site.

Under high soil moisture conditions, plants maintain open stomata to maximize carbon assimilation without risk of hydraulic stress, resulting in elevated transpiration rates. High transpiration reduces the residence time of water in leaves, thereby limiting evaporative δ^18^O enrichment and producing lower (more negative) δ^18^O values in cellulose (Barbour et al., 2004; Gessler et al., 2014; Cernusak et al., 2016). Conversely, during water-limited periods, stomatal closure reduces transpiration, prolonging leaf water residence time and enhancing evaporative enrichment, which results in higher δ^18^O values in tree-ring cellulose (Barbour et al., 2004; 2007). Overall, the combined isotope, eddy covariance, and modelling evidence indicates that carbon–water interactions at the Lägeren forest are governed by both physiological regulation and hydrological buffering. While isotope-derived _i_WUE and process-based simulations consistently captured long-term physiological adjustments, ecosystem-scale WUE reflected additional controls associated with ET partitioning, species composition, and ecosystem structure. Together, these findings demonstrate that interpreting forest water-use efficiency requires explicit consideration of the spatial and temporal scales represented by each measurement approach.

## 5. Conclusions

WUE is widely used to assess forest responses to climate change, yet its interpretation depends strongly on the spatial and temporal scales represented by different measurement approaches. Rather than providing alternative estimates of the same process, tree-ring stable isotopes, eddy covariance observations, and process-based models each capture complementary aspects of carbon–water interactions. Our study demonstrates that integrating these approaches provides a robust mechanistic framework for interpreting WUE across biological scales and for improving predictions of forest responses to increasing atmospheric CO_2_, rising atmospheric water demand, and future climatic extremes. Within this framework, the main findings of our study are:

- The strong agreement between _i_WUE^obs^ and _i_WUE^mod^ (*r* = 0.61, p < 0.05) is a direct consequence of the species-specific consistency between approaches: both the isotopic chronology and the 3D-CMCC-FEM model simulations represent exclusively the physiology of mature *F. sylvatica*, confirming that the model successfully captures leaf-level physiological coupling between photosynthetic assimilation and stomatal conductance for this species.
- The multi-isotope evidence demonstrates that the observed ∼20% increase in _i_WUE over the study period is predominantly driven by enhanced photosynthetic assimilation (*A*) under rising CO_2_, rather than by progressive stomatal closure, as source water dynamics rather than evaporative enrichment dominated the δ^18^O and δ²H signals at Lägeren.
- The divergence between WUE_eco_^obs^ and WUE_eco_^mod^ is primarily attributable to the systematic underestimation of ET by the model (∼50% of observed values), resulting from its single-species old-growth deciduous configuration that excludes the water use of co-occurring broadleaf species, evergreen conifers, younger trees, and understory vegetation, the latter contributing approximately 10% of above-canopy ET at Lägeren.
- Model–data agreement improved substantially with temporal aggregation, indicating that the model reproduces seasonal and interannual patterns more reliably than short-term variability. The improvement from daily to monthly timescales highlights the strong influence of high-frequency meteorological variability, atmospheric turbulence, and flux measurement uncertainty on ecosystem-scale WUE.
- The absence of a significant relationship between intrinsic and ecosystem WUE demonstrates that these metrics should not be interpreted interchangeably. Intrinsic WUE reflects leaf-level physiological regulation, whereas ecosystem WUE integrates additional hydrological, structural, and community-level processes. Consequently, long-term reconstructions of iWUE based on tree-ring δ^13^C cannot be directly extrapolated to ecosystem-scale carbon–water exchange without considering these scale-dependent processes

Overall, this study presents, to our knowledge, the first mechanistic framework integrating tree-ring stable isotopes, eddy covariance observations and process-based ecosystem modelling to disentangle WUE from leaves to ecosystems. This integrative approach moves beyond simple comparisons among methodologies by identifying the biological and hydrological processes responsible for scale-dependent carbon–water coupling, thereby providing a new framework for improving predictions of forest responses to global change.

## Acknowledgments

PFP was funded by two grants from COST Action CA21138 CLEANFOREST “Joint effects of Climate Extremes and Atmospheric Deposition on European FORESTS”, supported by COST (European Cooperation in Science and Technology). (1) a Short-Term Scientific Mission (STSM) at WSL, and (2) a Young Researcher and Innovator Conference Grant (YRICG). This work has been partially supported by MIUR Project (PRIN 2020) “Unraveling interactions between WATER and carbon cycles during drought and their impact on water resources and forest and grassland ecosySTEMs in the Mediterranean climate (WATERSTEM)” (Project num-ber: 20202WF53Z).

## Author contributions

PFP: Writing – review & editing, Writing – original draft, Visualization, Investigation, Formal analysis, Data curation, Conceptualization, Funding acquisition; DD: Writing – review & editing, Validation, Methodology, Formal analysis; DC: Writing – review & editing; EV: Methodology, Data curation; MML: Methodology, Formal analysis, Conceptualization, review & editing; MS: review & editing, Investigation, Formal analysis, Project administration, Funding acquisition; SK: review & editing, Formal analysis, Data curation, Conceptualization; LH: review & editing, Methodology, Formal analysis, Data curation; AC: Writing – review & editing, Writing – original draft, Project administration, Funding acquisition.

## Data availability

**Tree-ring chronology and triple isotopes data.** Data will be made available on request to the corresponding author.

**Modelled data are deposited and publicly available in Zenodo:** https://doi.org/10.5281/zenodo.20928067

**Eddy covariance (EC) observational data can be accessed from the following public repositories:** https://www.swissfluxnet.ethz.ch/index.php/sites/site-info-ch-lae/

The source code used in this study is freely available at the same Zenodo repository as the modelled data.

